# *MaTAR25* LncRNA Regulates the *Tensin1* Gene to Impact Breast Cancer Progression

**DOI:** 10.1101/2020.02.03.931881

**Authors:** Kung-Chi Chang, Sarah D. Diermeier, Allen T. Yu, Lily D. Brine, Suzanne Russo, Sonam Bhatia, Habeeb Alsudani, Karen Kostroff, Tawfiqul Bhuiya, Edi Brogi, C. Frank Bennett, Frank Rigo, David L. Spector

## Abstract

Misregulation of long non-coding RNA genes has been linked to a wide variety of cancer types. Here we report on <u>Ma</u>mmary <u>T</u>umor <u>A</u>ssociated <u>R</u>NA 25 (*MaTAR25*), a nuclear enriched and chromatin associated lncRNA that plays a role in mammary tumor cell proliferation, migration, and invasion, both *in vitro* and *in vivo*. *MaTAR25* functions by interacting with purine rich element binding protein B (PURB), and associating with a major downstream target gene *Tensin 1* (*Tns1*) to regulate its expression in *trans*. Knockout of *MaTAR25* results in down-regulation of *Tns1* leading to a reorganization of the actin cytoskeleton, and a reduction of focal adhesions and microvilli. The human ortholog of *MaTAR25*, *LINC01271*, is upregulated with human breast cancer stage and metastasis.

**SIGNIFICANCE:** LncRNAs have great potential to reveal new regulatory mechanisms of function as well as having exciting therapeutic capacity given their ease of being targeted by nucleic acid drugs. Our study of *MaTAR25,* and its human ortholog *LINC01271,* reveal an unexpected function of this lncRNA in breast cancer progression by regulating *Tns1* gene expression, whose protein product is a critical component of focal adhesions linking signaling between the extracellular matrix and the actin cytoskeleton. We identified *LINC01271 as* the human ortholog of *MaTAR25*, and importantly, increased expression of *LINC01271* is associated with poor patient prognosis and cancer metastasis. Our findings demonstrate that *LINC01271* represents an exciting therapeutic target to alter breast cancer progression.

## INTRODUCTION

Breast cancer is the most common cancer among women in the United States and world-wide, with an estimated 268,600 new cases of invasive disease in women in the United States in 2019 (1). Although breast cancer mortality has been decreasing over the past two decades, it is still the second leading cause of cancer deaths in American women accounting for 15% of all cancer deaths. Breast tumors can be classified into multiple subtypes based on histological evaluation and the most frequent type of breast tumors are ductal carcinomas, which affect the milk ducts of the breast. Ductal carcinomas can further be separated into two groups: non-invasive ductal carcinoma *in situ* (DCIS), and invasive ductal carcinoma (IDC) which accounts for 75% of all breast cancers (2, 3). Breast cancer is recognized as a heterogeneous disease and molecular classification of invasive breast carcinomas can stratify tumors into informative subtypes and provide key prognostic signatures. In addition to traditional pathological characterization and immunohistochemistry (IHC) to examine protein levels of markers such as estrogen receptor (ER), progesterone receptor (PR) and epidermal growth factor receptor-2 (HER2), additional studies evaluating genomic rearrangements and molecular expression profiles of breast cancers have provided further genetic insights to better understand the disease (4–6). These approaches have identified six major molecular subtypes of breast cancer (luminal A, luminal B, HER2-enriched, triple negative/basal-like, normal breast-like, and claudin-low) (7), each displaying different phenotypic and molecular features and which have distinct clinical outcomes.

In recent years, large scale genome-wide studies indicated that thousands of RNAs can be transcribed from the human and mouse genomes that lack protein-coding capacity (8–10). In particular, long non-coding RNAs (lncRNAs) with a length *≥* 200 nucleotides have been suggested to play key roles in a diverse range of biological processes (11–13). Most lncRNAs are capped, spliced, and poly-adenylated (8). In addition, many lncRNAs are expressed in a tissue-specific and/or cell type specific manner, and are involved in various gene regulatory pathways (14, 15). Furthermore, misregulation of lncRNA expression has been linked to various diseases including neuromuscular diseases, developmental disorders, neurodegenerative diseases, and cancers (16–20). Several lncRNAs have been implicated as regulatory molecules in breast cancer progression and metastasis through different mechanisms (21, 22). For example, the *HOX* antisense intergenic RNA (*HOTAIR*) is overexpressed in primary breast tumors and can alter the localization pattern of Polycomb repressive complex 2 (PRC2) and histone methylation to regulate gene expression in breast carcinoma cells impacting breast cancer progression and metastasis (23). Recent findings suggest that the lncRNA breast cancer anti-estrogen resistance 4 (*BCAR4*) (24) can control GLI family zinc finger 2 (GLI2) gene expression to promote cancer cell migration by interacting with Smad nuclear interacting protein 1 (SNIP1) and serine/threonine-protein phosphatase 1 regulatory subunit 10 (PNUTS). Targeting *BCAR4* by locked nucleic acids (LNA) in mouse models significantly affects cancer cell invasion and reduces lung metastases (25). Genetic knockout or ASO-mediated knockdown of *<u>M</u>etastasis <u>A</u>ssociated <u>L</u>ung <u>A</u>denocarcinoma <u>T</u>ranscript <u>1</u> (Malat1)* was shown to result in differentiation of primary mammary tumors and a significant reduction in metastasis (26). In addition to transcriptional regulation, lncRNAs can have other regulatory roles. For example, the lncRNA *PVT1* has been shown to stabilize the Myc oncoprotein in breast cancer cells (27), and the lncRNA NKILA can interact with and stabilize the NF-κB/IκB complex and inhibit breast cancer metastasis (28). However, for the majority of lncRNAs, the exact function and molecular mechanism of action in breast cancers still awaits detailed characterization. Previously, we performed an RNA sequencing (RNA-seq) screen to identify differentially expressed lncRNAs between mammary tumor cells and normal mammary epithelial cells. From this screen, we identified 30 previously uncharacterized lncRNAs as <u>Ma</u>mmary <u>T</u>umor <u>A</u>ssociated <u>R</u>NAs (*MaTARs*) 1-30 (29).

Here, we examined the role of *MaTAR25* in mammary tumor progression and metastasis. We find that genetic knockout of *MaTAR25* in highly aggressive 4T1 triple negative (ER-, PR-, HER2-) mammary carcinoma cells results in a reduction in cell proliferation, migration, and invasion. Knockout cells transplanted into the mammary fat pad of BALB/c mice results in a significant decrease in tumor growth as compared to 4T1 control cells. Further, tail vein injection of luciferase labeled *MaTAR25* knockout cells showed reduced homing to the lungs and a significant decrease in metastatic nodules. In a complementary study, antisense oligonucleotide (ASO) mediated knockdown (KD) of *MaTAR25* in the MMTV-Neu-NDL mouse model resulted in a significant decrease in tumor growth and a reduction in lung metastases. Analysis of the molecular function of *MaTAR25* indicates that it regulates the *Tns1* gene at the transcriptional level. Loss of *MaTAR25* results in a reduction of *Tns1* at the RNA and protein levels and a subsequent reorganization of the actin cytoskeleton and a reduction in focal adhesions and microvilli. Together, our data reveal *MaTAR25*, and its identified human ortholog *LINC01271*, as an exciting therapeutic candidate whose expression can be altered to impede breast cancer progression and metastasis.

## RESULTS

### Characterization of *MaTAR25*, a nuclear enriched lncRNA

We previously performed an RNA-seq screen to identify lncRNAs over-expressed in mammary tumors vs normal mammary epithelial cells as a means to identify potential candidates involved in mammary cancer progression, and to explore their potential as therapeutic targets or key biomarkers in human breast cancer (29). Among those lncRNA genes identified, the *MaTAR25* gene on mouse chromosome 2 was originally annotated as 1200007C13Rik and it encodes a single transcript containing two exons (Fig. 1A). *MaTAR25* is overexpressed in mammary tumors in the MMTV-Neu-NDL (HER2 subtype) model compared to normal mammary epithelial cells and it is also upregulated in luminal and triple negative sub-types of mammary cancer (29). Analysis of ENCODE and FANTOM5 RNA-seq data has shown there is little to no expression of *MaTAR25* in normal mouse tissues compared to MMTV-Neu-NDL tumor cells (29, 30) (Supplementary Fig. S1A). The full length *MaTAR25* transcript was determined to be 1,978 nucleotides by 5’ and 3′ rapid amplification of cDNA ends (RACE) and Sanger sequencing (Fig. 1B), which was further confirmed by Northern blot analysis (Fig. 1C).

**Figure 1.**
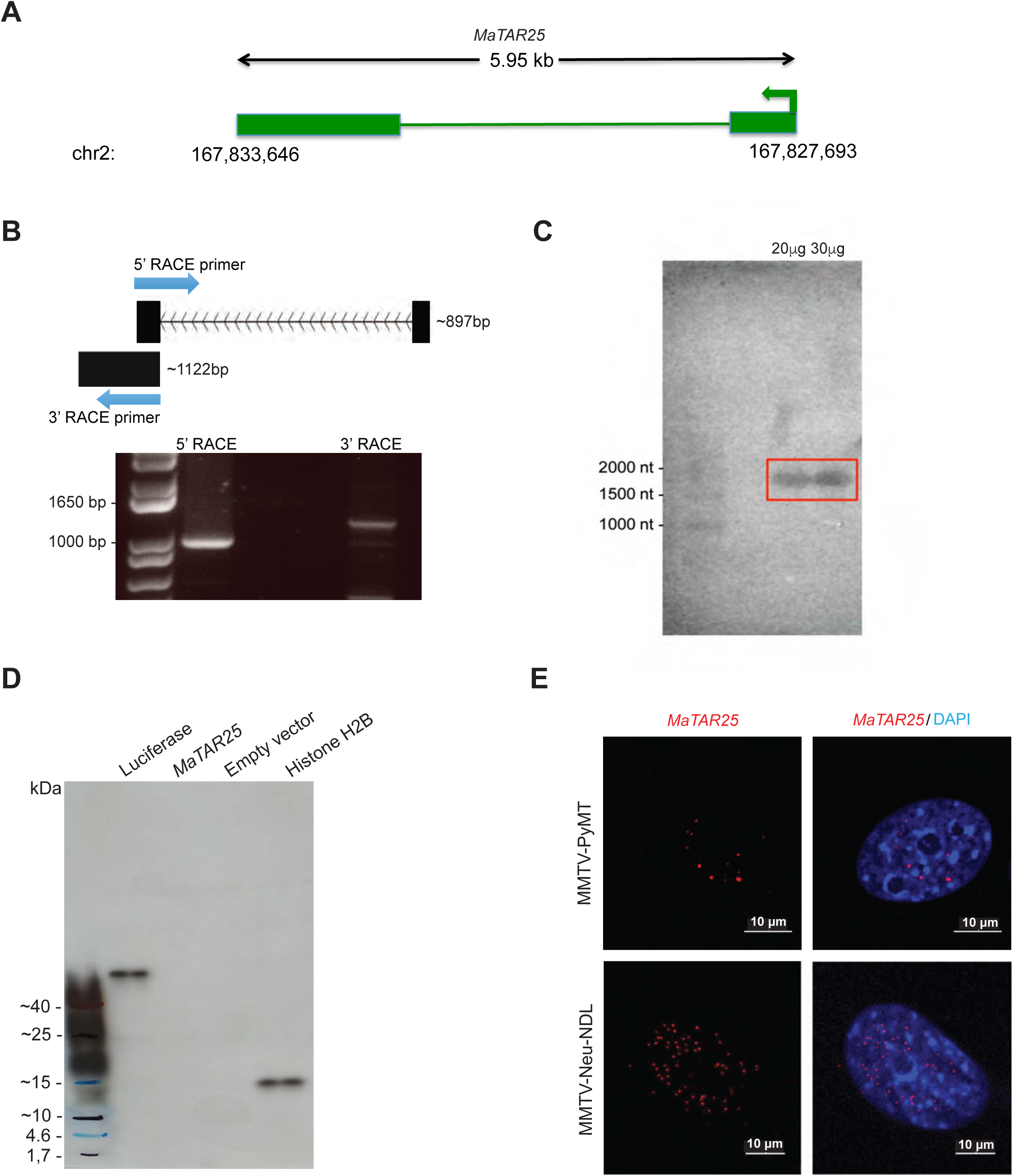
Characterization of Mammary Tumor Associated RNA 25 (MaTAR25) (A) Representation of the *MaTAR25* gene locus. *MaTAR25* is an intergenic lncRNA gene located on mouse chromosome 2, and the *MaTAR25* RNA transcript contains 2 exons and a poly (A) tail. (B) 5’ and 3’ rapid amplification of cDNA ends (RACE) was performed to identify the full length *MaTAR25* transcript. (C) The full length *MaTAR25* transcript was confirmed by Northern blot analysis to be ∼2000 nt. 20 μg or 30 μg total RNA extracted from MMTV-PyMT primary cells was electrophoresed on a 1% agarose gel and probed. (D) *In vitro* transcription and translation reactions were performed to confirm that *MaTAR25* does not produce a peptide. The reaction products were run on a 4-20% gradient SDS-PAGE gel, and the signals were detected by HRP-conjugated Streptavidin. Luciferase control DNA and *Xenopus laevis* Histone H2B (HISTH2B) expressing plasmids were used as positive controls and empty vector as a negative control. (E) Representative smRNA-FISH images showing localization of *MaTAR25* RNA transcripts within nuclei of MMTV-PyMT and MMTV-Neu-NDL primary cells. Scale bars are 10 μm.

According to three independent computational coding potential prediction programs, the *MaTAR25* RNA transcript has very low protein coding potential and is suggested to be a non-coding RNA (Supplementary Fig. S1B-S1D). However, there is one predicted open reading frame (ORF) with the potential to generate a 123 amino acid peptide (∼13 kDa). In order to assess whether a peptide is encoded by the *MaTAR25* transcript we performed *in vitro* transcription and translation. Compared to a luciferase DNA control (expected size 61 kDa) and a *Xenopus laevis* Histone H2B (HISTH2B) expressing plasmid control (expected size 14 kDa), there was no detectable peptide generated from a plasmid that contained the *MaTAR25* sequence (Fig. 1D). Together, these computational and experimental results confirm that *MaTAR25* does not make a peptide, and thus is a bona fide lncRNA.

In order to determine the localization and abundance of *MaTAR25* we performed single molecule RNA fluorescence in situ hybridization (smRNA-FISH) to detect *MaTAR25* RNA transcripts within MMTV-PyMT (luminal B) and MMTV-Neu-NDL (Her2/neu+) primary mammary tumor cells. The majority of *MaTAR25* transcripts were detected in cell nuclei (Fig. 1E) and each nucleus contained ∼10-15 transcript foci. Thus, *MaTAR25* is a nuclear-enriched lncRNA with a potential role in the regulation of gene expression in mammary cancer cells.

### *MaTAR25* knockout decreases 4T1 cell viability/migration/invasion

To assess the functional role of *MaTAR25*, we proceeded to genetically knockout (KO) *MaTAR25* in highly aggressive 4T1 triple negative (ER-, PR-, HER2-) mammary carcinoma cells using CRISPR/Cas9. We designed gRNA pairs targeting various regions upstream and downstream of the transcription start site (TSS) of *MaTAR25* to create a genomic deletion (Fig. 2A and Supplementary Fig. S2A). *MaTAR25* knockout clones were single cell sorted and selected by Sanger sequencing, qRT-PCR, as well as smRNA-FISH (Fig. 2A and Supplementary Fig. S2B).

**Figure 2.**
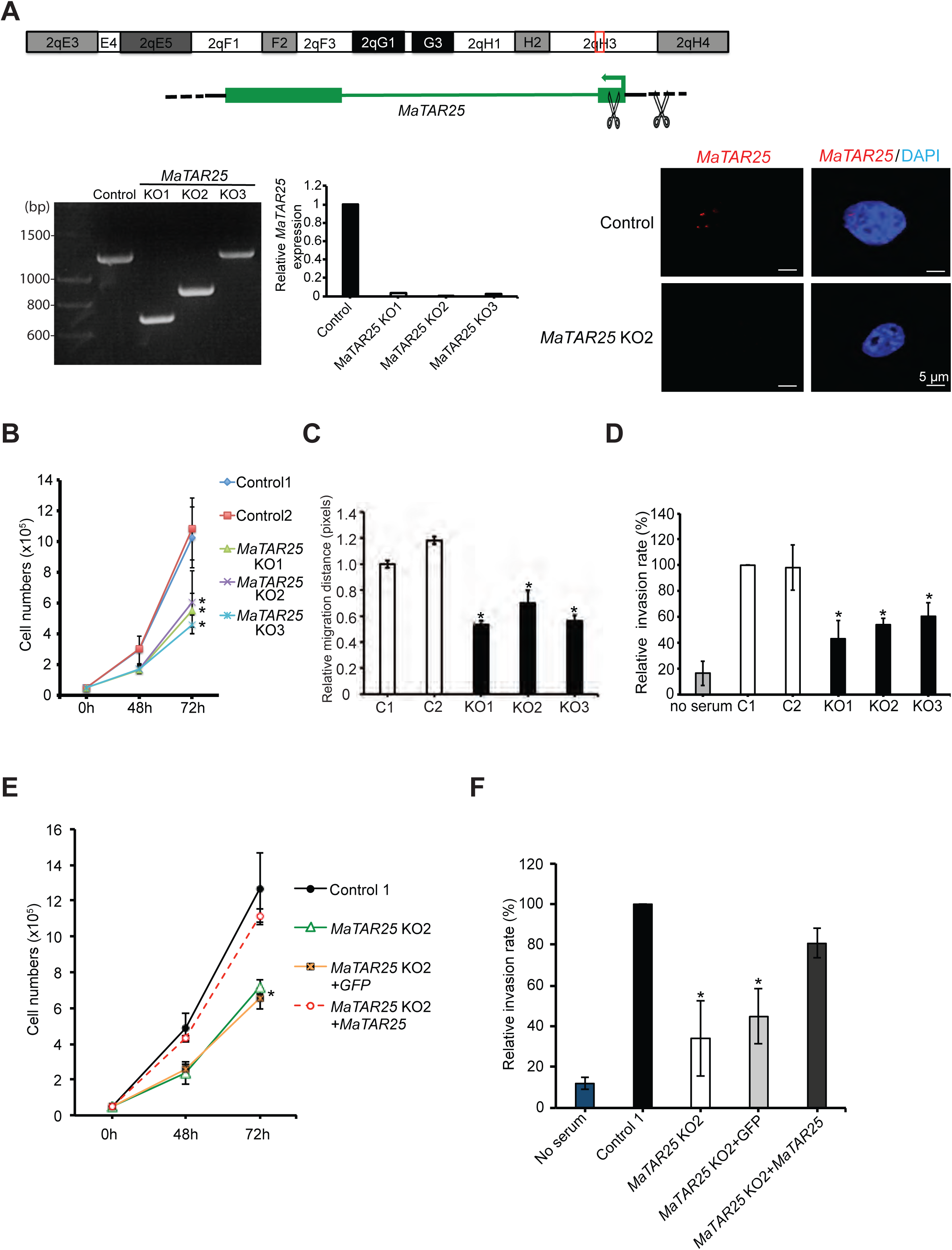
*MaTAR25* knockout affects 4T1 cell viability, migration, and invasion *in vitro*; all of which can be rescued by ectopic expression of *MaTAR25* in knockout cells. (A) CRISPR/Cas9 was used to generate *MaTAR25* KO clones in 4T1 cells. Pairs of sgRNAs were introduced targeting upstream and downstream of the transcription start site of *MaTAR25*, resulting in 390-620 bp genomic delections, and a *Renilla* Luciferase sgRNA was used as a negative control. Knockout clones were selected by genomic PCR and Sanger sequencing for homozygous genomic deletion. qRT-PCR and representative images of smRNA-FISH are shown to confirm *MaTAR25* KO. Scale bars are 5 μm. (B) 4T1 cells were seeded at the same cell density in 12-well tissue culture plates at day 0 and cell counting was performed at different time points. The mean cell numbers of three independent replicates of 4T1 control groups and *MaTAR25* KO groups is shown ± SD (n=3). \**p* < 0.05 (student’s t-test). (C) Live cell tracking was performed over time to examine cell migration. Images were collected every 5 minutes for a total of 8 hours and analyzed by CellTracker image processing software. The mean relative migration distance (μm) of two independent replicates of 4T1 control groups and *MaTAR25* KO groups is shown ± SD (n=3). \**p* < 0.05 (D) 24-well Boyden chamber invasion assay (24 hours). The mean relative cell invasion of three independent replicates of 4T1 control groups and *MaTAR25* KO groups is shown ± SD (n=3). \**p* < 0.05 (student’s t-test). (E) Ectopic expression of *MaTAR25* or GFP was used as positive and negative controls to assess rescue in a cell viability assay, or (F) cell invasion assay. The mean cell numbers and mean relative cell invasion of three independent replicates of 4T1 Control1, *MaTAR25* KO2, *MaTAR25* KO2 with GFP expression, and *MaTAR25* KO2 with *MaTAR25* ectopic expression is shown ± SD (n=3) \**p* < 0.05 (student’s t-test).

After selecting several *MaTAR25* KO clones, we evaluated them for alterations in cell viability, migration, and invasion as compared to 4T1 control cells. *MaTAR25* KO cells exhibited a significant decrease of 50% in cell viability as compared to 4T1 control cells (Fig. 2B). To further investigate this phenotype, we performed BrdU labeling and FACS analysis, which demonstrated a two-fold increase in G2 cells suggesting that the decreased proliferation phenotype is most likely the result of a lengthened G2 phase (Supplementary Fig. S2C). As cell migration and invasion are critical processes associated with metastasis we were interested in determining whether *MaTAR25* loss might play a role in these events. We used a live cell tracking assay to assess cell migration and we found a 40% reduction in cell motility upon loss of *MaTAR25* (Fig. 2C and Supplementary Fig. S2D). A wound healing assay also corroborated the observed difference in cell migration between 4T1 control and *MaTAR25* KO cells (Supplementary Fig. S2E). Finally, we used a Boyden chamber invasion assay and found that loss of *MaTAR25* resulted in a 45% reduction in invasion ability as compared to 4T1 control cells (Fig. 2D).

In order to exclude the possibility that the phenotypes observed in *MaTAR25* KO cells were caused by disrupting chromatin structure rather than specific loss of the *MaTAR25* transcript, we generated single cell ectopic overexpression clones of *MaTAR25* in 4T1 *MaTAR25* KO cells. Ectopic expression of *MaTAR25* rescued both the cell proliferation and invasion phenotypes (Fig. 2E-2F), indicating that *MaTAR25* RNA plays an important role in these processes *in situ,* and likely exhibits its effect *in trans*. Hence, *MaTAR25* appears to be an important lncRNA impacting mammary tumor cell growth and critical aspects of metastasis. To further explore *MaTAR25’s* downstream targets, we performed RNA-seq to identify differentially expressed genes by comparing *MaTAR25* KO cells with 4T1 control cells (Supplementary Table S1). Pathway analysis of the differentially expressed genes by Kyoto Encyclopedia of Genes and Genomes (KEGG) and Gene Set Enrichment Analysis (GSEA) revealed alteration in cell cycle and DNA related processes, both related to the phenotypes we observed in *MaTAR25* KO cells (Supplementary Fig. S2F).

### *MaTAR25* knockout decreases tumor progression/metastasis *in vivo*

In order to further evaluate the functional impact and the therapeutic potential of *MaTAR25* in mammary tumor progression, we performed multiple *in vivo* studies. Injection of *MaTAR25* 4T1 KO cells into the mammary fat pad of BALB/c mice resulted in a significant 56% decrease in tumor growth at day 28, compared to the 4T1 control injected group (Fig. 3A-3B). In addition, we performed tail vein injection using *MaTAR25* KO cells expressing a luciferase reporter to track cancer cell homing and metastasis to the lungs in BALB/c mice. The *in vivo* bioluminescence signal in the lungs of mice injected with *MaTAR25* KO cells was reduced (Supplementary Fig. S3A) compared to those injected with 4T1 control cells. At day 21, the number of metastatic nodules in lung samples collected from the *MaTAR25* KO group was also significantly decreased by 62% compared to the 4T1 control group (Fig. 3C).

**Figure 3.**
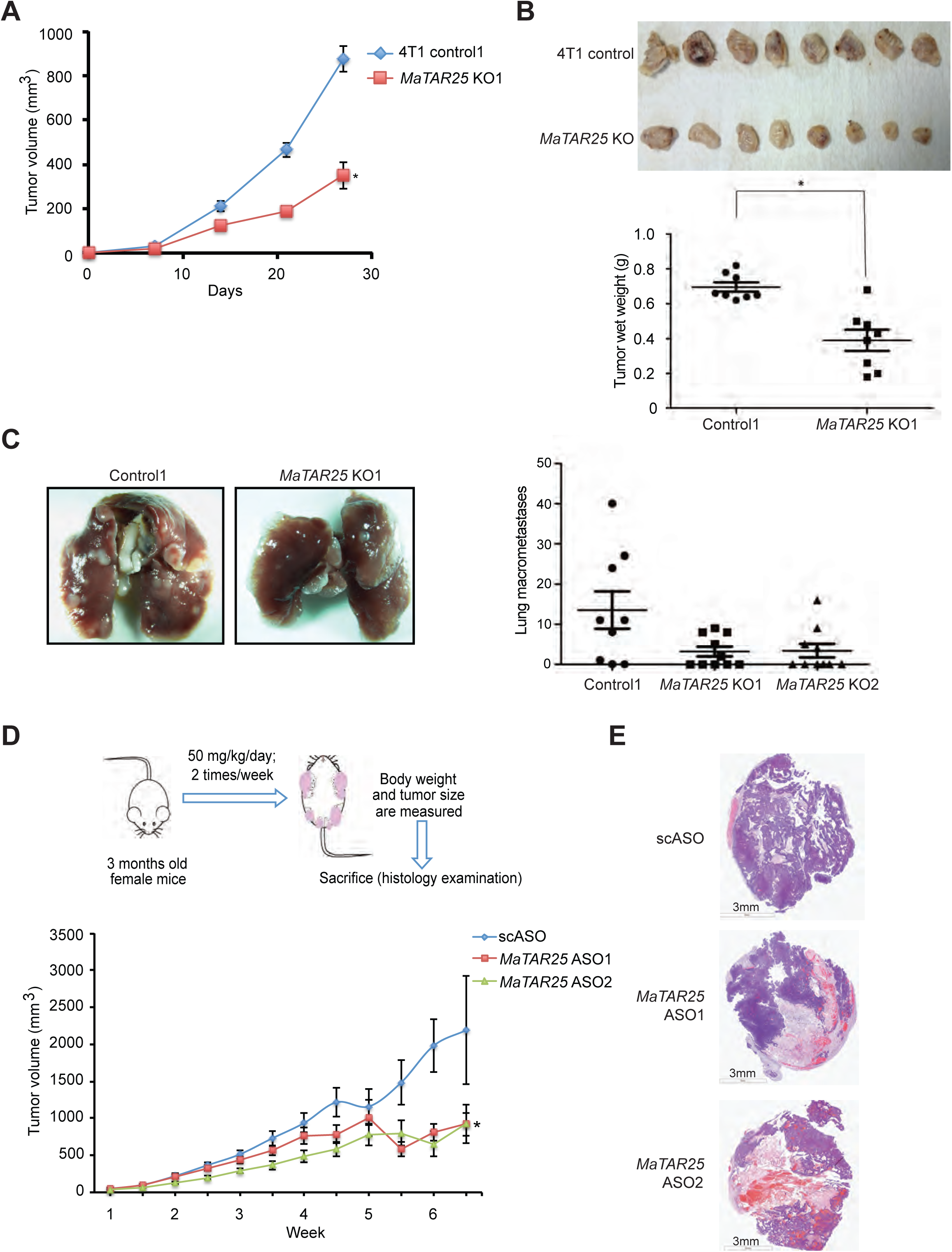
*MaTAR25* knockout impairs tumor growth and metastasis *in vivo*. (A) 4T1 control or *MaTAR25* KO cells were injected orthotopically into the mammary fat pad of female BALB/c mice. Primary tumors were measured every week over a period of four weeks and the mean tumor volume of 8 mice per group is shown ± SE. \**p* < 0.05 (student’s t-test). (B) Mice were sacrificed and tumors were collected at day 28 to compare the tumor growth rate between the control group and *MaTAR25* KO groups. Tumors derived from *MaTAR25* KO cells showed a 56% reduction in tumor growth. The mean tumor wet weight is shown ± SE. \**p* < 0.05 (student’s t-test). (C) Female BALB/c mice were injected into the tail vein with 4T1 Control1 or *MaTAR25* KO cells. Mice were monitored every week and sacrificed at day 21. Mouse lungs were collected and imaged (left panel), and lung metastatic nodules were counted to compare the metastatic ability between the control group and *MaTAR25* KO group (right panel). Mice injected with *MaTAR25* KO cells exhibited a 62% reduction in lung metastatic nodules. (D) Schematic showing the approach for ASO mediated knockdown of *MaTAR25* in MMTV-Neu-NDL mice. Two independent *MaTAR25* ASOs or a control scASO were used for subcutaneous injection. Primary tumors were measured twice per week and the mean tumor volume of 7 mice per group is shown ± SE. \**p* < 0.05 (student’s t-test). (E) Representative hematoxylin and eosin (H&E) stained tumor images showing the different histological phenotypes between tumor samples from scASO injected mice and *MaTAR25* ASO injected groups.

As a complementary approach to CRISPR/Cas9 KO we designed a series of *MaTAR25* specific antisense oligonucleotides (ASOs), (16mers) comprised of phosphorothioate-modified short S-cEt (S-2 ′ -O-Et-2′, 4′ -bridged nucleic acid) gapmer chemistry (31–33). We individually screened multiple ASOs targeting *MaTAR25* to identify the most effective ASOs in terms of knockdown (KD) efficiency by qRT-PCR after 48 hours and 72 hours of ASO treatment in 4T1 cells. The two most effective *MaTAR25* ASOs achieved a knockdown ranging from 70-90% (Supplementary Fig. S1E). When comparing *MaTAR25* ASO treated cells to mock or scrambled ASO (scASO) treated 4T1 control cells after 72 hours, we found a significant decrease in cell viability using cell counting assays (-45% for ASO1 and -38% for ASO2) (Supplementary Fig. S1F), consistent with our KO studies indicating that *MaTAR25* has a role in mammary cancer cell proliferation.

Furthermore, to assess the therapeutic potential of reducing the level of *MaTAR25 in vivo*, we evaluated the impact of subcutaneous injection of two independent *MaTAR25* ASOs for their *in vivo* ability to knockdown *MaTAR25* and to impact mammary tumor progression in the MMTV-Neu-NDL mouse model. ASO mediated knockdown of *MaTAR25* resulted in a 59% decrease in tumor growth compared to the scASO control group (Fig. 3D and Supplementary Fig. S3B). By comparing the hematoxylin and eosin (H&E) stained tumor sections collected from the *MaTAR25* ASO injected group with the scASO control group, we observed a strong level of necrosis in the *MaTAR25* ASO treated mammary tumor samples (Fig. 3E) but not in other non-tumor tissues (Supplementary Fig. S3C). Importantly, mammary tumors from the scASO control group lacked any significant necrotic phenotype (Fig. 3E). We also collected lung samples from each group to examine for the presence of micro-metastases, and the H&E stained lung sections showed that KD of *MaTAR25* resulted in a 40% incidence rate of micro-metastatic nodules in lungs from ASO1 or ASO2 treated animals as compared to a 76.9% incidence rate for the scASO control group (Supplementary Fig. S3D). Together, our *in vitro* and *in vivo* data indicate that *MaTAR25* plays a critical role in promoting mammary tumor progression and metastasis.

### *MaTAR25* is a positive upstream regulator of *Tns1*, a mediator of cell-matrix adhesion and migration

Next, we were interested in revealing aspects of the molecular mechanism of action of *MaTAR25* in regulating mammary tumor progression. Since we previously identified *MaTAR25* to be highly enriched in cell nuclei by smRNA-FISH we went on to perform cell fractionation to isolate cytoplasmic and nucleoplasmic lysates as well as chromatin pellets of 4T1 cells to determine the subcellular enrichment of *MaTAR25* by qRT-PCR analysis. Notably, compared to the enrichment of β*-actin* and *Malat1*, we found a significant enrichment of *MaTAR25* in the nucleoplasmic and chromatin fractions (Fig. 4A), indicating that the molecular mechanism of action of *MaTAR25* may be related to transcriptional regulation. To test this hypothesis, we performed <u>Ch</u>romatin Isolation by <u>R</u>NA <u>P</u>urification (ChIRP) (34) to pull down RNA/DNA complexes by using specific biotin-labeled antisense oligonucleotides targeting *MaTAR25* (Fig. 4B) as well as biotin-labeled antisense oligonucleotides targeting housekeeping gene PPIB transcripts as the corresponding control. ChIRP-seq identified *MaTAR25* genomic targeting sites, and revealed that these targets are highly enriched in simple repeats regions and LTRs (log ratio enrichments to input are 2-3 fold) (Supplementary Fig. S4A). According to motif analysis these regions are potential binding sites of the transcription factors ZNF354C, TEAD, GATA1, and REL (data not shown). Combining the *MaTAR25* KO RNA-seq data and ChIRP-seq results, we found a total of 446 overlapping genes (Fig. 4C and Supplementary Table S1), which could be downstream targets regulated by *MaTAR25*. Among these overlapping genes, the top gene ranked by ChIRP-seq data, just under *MaTAR25* itself, is *Tensin1* (*Tns1*) (Fig. 4C and Supplementary Fig. S4B). The *Tns1* gene encodes for a protein that localizes to focal adhesions and positively regulates cell migration and invasion (35, 36). By qRT-PCR and immunoblot analysis, we found that the RNA and protein levels of *Tns1* are significantly lower in *MaTAR25* KO cells than in 4T1 control cells (Fig. 4D). Interestingly, ectopic expression of *MaTAR25* in *MaTAR25* KO cells results in a correspondingly increased level of *Tns1* (Fig. 4E). Hence, we conclude that *Tns1* is a direct downstream target of *MaTAR25* and further confirming that it imparts its function *in trans*. In order to confirm the ChIRP-seq result and to further investigate how *MaTAR25* might regulate the level of *Tns1*, we next performed double-label DNA-FISH to detect the *MaTAR25* (Chr2) and *Tns1* (Chr1) gene loci in cells, and we found no physical interaction between these genomic loci (Supplementary Fig. S4D, upper panels). However, combined *MaTAR25* smRNA-FISH and *Tns1* DNA-FISH in the same cells showed that *MaTAR25* RNA was overlapping with at least one *Tns1* allele in 50% of the cells (Supplementary Fig. S4D, lower panels). This suggests that *MaTAR25* RNA transcripts can bind to the gene body of *Tns1* to regulate its expression. We therefore performed CRISPR/Cas9 knockout using gRNAs targeting *Tns1* in 4T1 cells and selected *Tns1* KO clones for *in vitro* functional assays. We found that the *Tns1* KO cells phenocopied the *MaTAR25* KO cells and exhibited a significant 40% decrease in cell viability (Fig. 4F) and a 30% decrease in cell migration vs control cells (Supplementary Fig. S4E). In addition, ectopic expression of *Tns1* in 4T1 *MaTAR25* KO cells can rescue the cell viability phenotype (Fig. 4G). Interestingly, high expression of *TNS1* is strongly correlated with poor survival of grade 3 breast cancer patients (37) (Supplementary Fig. S4C). Together, these data indicate that *Tns1* is a critical downstream target of *MaTAR25*.

**Figure 4.**
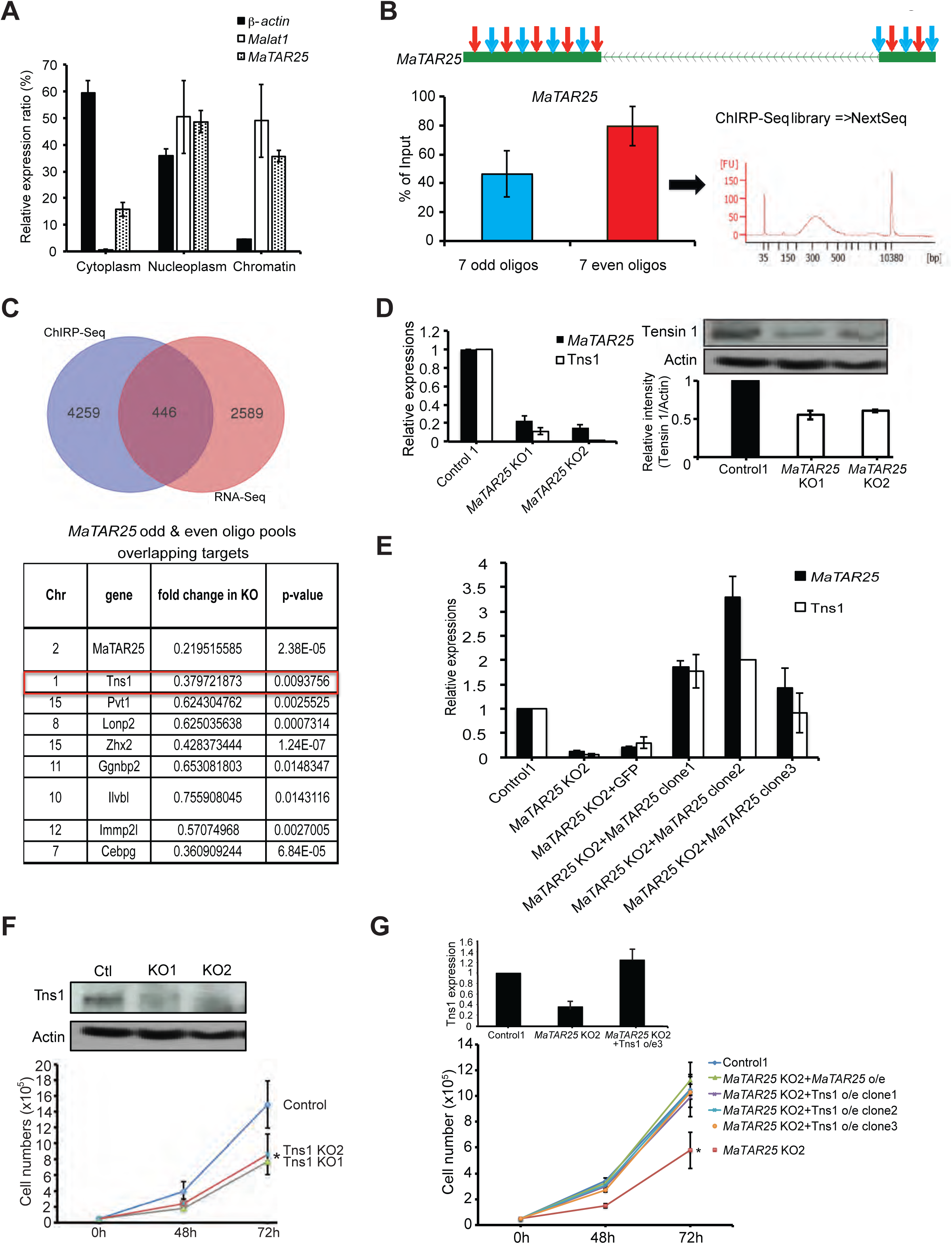
*MaTAR25* is a positive upstream regulator of *Tns1*. (A) Cell fractionation was performed to isolate cytoplasmic, nucleoplasmic, and chromatin associated RNA. qRT-PCR was used to determine the subcellular localization ratio of *MaTAR25* transcripts. *β-actin* and *Malat1* were used as marker RNAs for quality control of cell fractionation. (B) Schematic diagram showing the targeting of biotin labeled oligonucleotides binding *MaTAR25* transcripts for chromatin isolation by RNA purification (ChIRP)-seq. Odd and even oligo pools (7 oligos binding different regions within the *MaTAR25* transcript in each pool) were used for ChIRP-seq, and qRT-PCR was performed to assess RNA purification enrichment. (C) Venn diagram showing differentially expressed genes in *MaTAR25* KO cells identified from RNA-Seq overlapped with *MaTAR25* ChIRP-seq data. A total 446 overlapping genes were identified, and the top candidate genes are listed. (D) Validation of *Tns1* as a *MaTAR25* targeted gene by qRT-PCR and immunoblotting in 4T1 control and *MaTAR25* KO cells. (E) The RNA expression level of *Tns1* is rescued upon ectopic expression of *MaTAR25* in *MaTAR25* KO cells as determined by qRT-PCR. (F) CRISPR/Cas9 targeting was used in 4T1 cells to generate *Tns1* knockout clones. The upper panel shows expression levels of *Tns1* in 4T1 control, *Tns1* KO Clone1, and *Tns1* KO Clone2 by immunoblotting. The lower panel shows the cell counting viability assay results of three independent replicates of 4T1 Control1, *Tns1* KO1, and *Tns1* KO2. Results are mean ± SD (n=3) \**p* < 0.05 (student’s t-test). (G)Ectopic expression of *Tns1* in *MaTAR25* KO cells rescues the cell viability defect. The top panel shows expression levels of *Tns1* in 4T1 control1, *MaTAR25* KO2, *MaTAR25* KO2 with *Tns1* ectopic expression Clone1. The bottom panel shows the cell counting viability assay results of three independent replicates of 4T1 Control1, *MaTAR25* KO2, *MaTAR25* KO2 with *MaTAR25* ectopic expression, and *MaTAR25* KO with *Tns1* ectopic expression Clone1-3. Results are mean ± SD (n=3) \**p* < 0.05 (student’s t-test).

Since *Tns1* is a key component of focal adhesion complexes and is responsible for cell-cell and cell-matrix interactions as well as cell migration by interacting with actin filaments (38), we examined the organization of actin filaments, as well as the additional focal adhesion complex components paxillin and vinculin (39), in 4T1 control and *MaTAR25* KO cells by immunofluorescence (IF) confocal microscopy. Indeed, the F-actin microfilaments are disrupted (Fig. S4G1 and Supplementary Fig. S4G2) and the distribution of paxillin and vinculin proteins are altered dramatically (Supplementary Fig. S4H) in 4T1 *MaTAR25* KO cells as compared to 4T1 control cells. Interestingly, both ectopic expression of *MaTAR25* or *Tns1* in 4T1 *MaTAR25* KO cells can rescue the actin filament phenotype (Supplementary Fig. S4G3 and S4G4), supporting our finding that *Tns1* is a critical downstream target of *MaTAR25* regulating mammary tumor progression. To further evaluate the phenotype of *MaTAR25* KO cells we used transmission electron microscopy (TEM) (Supplementary Fig. S4F). TEM clearly revealed a dramatic 81% reduction of microvilli over the cell surface of *MaTAR25* KO cells compared to 4T1 control cells, indicating loss of *MaTAR25* expression impacts the actin bundling process (Supplementary Fig. S4H) as well as microvilli formation/maintenance in 4T1 cells.

### *MaTAR25* interacts with PURB to carry out its function

It has been suggested that lncRNAs can interact with transcriptional regulators/co-factors to form ribonucleoprotein (RNP) complexes to regulate the expression of downstream genes in the cell nucleus (40). To identify *MaTAR25* interacting proteins we used two different paired sets of biotin-labeled antisense oligonucleotides targeting *MaTAR25* for native RNA antisense oligonucleotide pull-down (RAP) in 4T1 cells followed by qRT-PCR which revealed a 50-60% pull-down efficiency (Supplementary Fig. S5A). Samples were eluted from beads for mass spectrometry isobaric tags for relative and absolute quantitative (MS-iTRAQ) analysis to identify proteins that bind to *MaTAR25*, and PPIB as the corresponding control. We ranked the candidate interactors based on detectable peptides above background in both pair sets of oligonucleotide pull-downs, and selected candidates with at least 2-fold enrichment compared to corresponding PPIB oligo pull-down (Fig. 5A). Among the protein candidates, two transcription co-regulators always appeared on the top list between multiple runs. These are purine rich element binding protein A (PURA) and purine rich element binding protein B (PURB), which can form homodimers or heterodimers in the nucleus (41). Additionally, one other protein, Y-box protein 1 (YBX1) also on the candidate list, but which did not pass the enrichment criteria, was shown in a previous study to interact with PURA to form a PURB/PURA/YBX1 heterotrimer (42). To verify our MS result, we first performed immunoblot analysis with PURA and PURB antibodies and we could detect extremely higher signals of PURA and PURB in samples of *MaTAR25* oligonucleotide pull-down than the PPIB oligonucleotide pull-down (Fig. 5B). When RAP was carried out with the same sets of oligonucleotide pairs using NDL cells, the MS result showed a greater enrichment of PURB than PURA (data not shown). Based on these data, we hypothesized that PURB is the lead protein directly binding to *MaTAR25*. Immunoblot analysis using pull-down samples from 4T1 and 4T1 *MaTAR25* KO cell lysates (Fig. 5B) plus pull-down samples from NDL primary cell lysates (Supplementary Fig. S5B) confirmed the specific interaction between *MaTAR25* and PURB. RNA immunoprecipitation (RIP) using PURB antibodies compared to IgG control also revealed the specificity of the *MaTAR25*-PURB interaction (Fig. 5C). To further confirm the role of PURB in regulating *Tns1* we manipulated the level of PURB in 4T1 cells either through ectopic overexpression (Fig. 5D) or siRNA mediated knockdown and demonstrated upregulation or down-regulation of *Tns1*, respectively (Supplementary Fig. S5C). Next, to determine if the interaction between *MaTAR25* and PURB is related to the expression of *Tns1* we examined the level of *Tns1* expression in each group. The results confirmed that the expression level of *Tns1* is related to the changes in PURB expression level in 4T1 cells, indicating that the *MaTAR25*/PURB RNP complex is essential for the regulation of *Tns1*.

**Figure 5.**
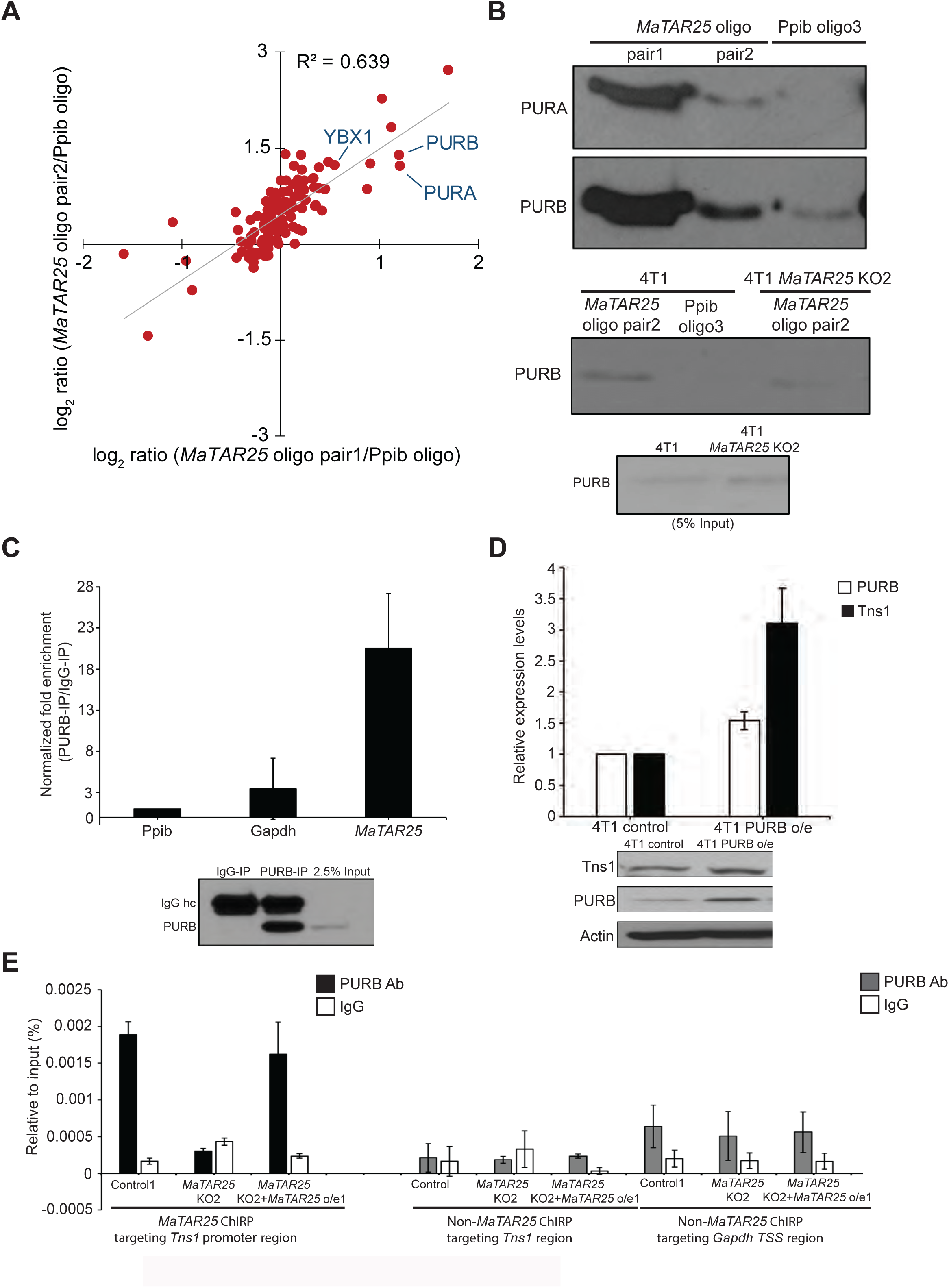
*MaTAR25* interacts with PURB to carry out its function. (A) Scatterplot depicts the fold enrichment of protein candidates from isobaric tags for the relative and absolute quantitation (iTRAQ) analysis comparing two independent oligo pair sets targeting *MaTAR25* RNA transcripts vs PPIB RNA transcripts. (B) Upper: immunoblot analysis of PURA and PURB following pull-down of *MaTAR25* or PPIB from 4T1 cells. Lower: immunoblot analysis of PURB following the pull-down of *MaTAR25* or PPIB from 4T1 cells or 4T1 *MaTAR25* KO cells. (C) *MaTAR25*, PPIB, and *Gapdh* transcripts were assessed by qRT-PCR in endogenous PURB, or IgG (negative control) immunoprecipitates from 4T1 cells. Fold enrichment over IgG signal is shown ± SD (n=3). Immunoblot analysis of PURB was performed as a control. (D) qRT-PCR analysis and immunoblotting of *Tns1* expression in 4T1 cells following ectopic over-expression of PURB. The relative expression levels are shown ± SD (n=3). (E) ChIP-qPCR analysis of PURB occupancy over the identified *MaTAR25* targeting region and non-targeting region of the *Tns1* DNA locus by ChIRP-seq analysis. ChIP-qPCR was performed in 4T1 control cells, 4T1 *MaTAR25* KO cells, and upon ectopic expression of *MaTAR25* in *MaTAR25* KO cells. Primers for a *MaTAR25* non-targeting region and the *Gapdh* TSS were used as negative controls. Bar graphs represent the mean ± SD (n=2).

*Tns1* isoform 3 was identified as the major isoform expressed in MMTV-Neu-NDL and 4T1 cells (data not shown). Based on our ChIRP-seq result, we were able to go on to identify the promoter region of *Tns1* isoform 3, which contains a very high purine:pyrimidine ratio (3:1) including many potential PURB binding sequence motifs (GGTGG) (43), as the main targeting region in the *Tns1* gene (Supplementary Fig. S5D). Moreover, Hypergeometric Optimization of Motif EnRichment (HOMER) motif analysis based on ChIRP-seq data indicated the top enriched motif sequence of *MaTAR25* interacting genes is GGTGGTGGAGAT further supporting the *MaTAR25* PURB binding motif sequence (Supplementary Fig. S5E). Therefore, we performed Chromatin immunoprecipitation (ChIP) using a PURB antibody and multiple qPCR primer pairs and showed that PURB has a high occupancy capacity over this region of *Tns1*. Importantly, the occupancy was impaired in *MaTAR25* KO cells and was able to be restored upon ectopic expression of *MaTAR25* in 4T1 *MaTAR25* KO cells (Fig. 5E). Together, these results provide compelling evidence indicating the interaction of PURB protein with *MaTAR25* is required for PURB binding to regulatory motifs in the *Tns1* gene in 4T1 cells.

### Human lncRNA *LINC01271* is the human ortholog of *MaTAR25*

In order to translate our exciting findings in regard to mouse *MaTAR25* to the human system for potential future clinical applications, we went on to characterize the human ortholog of *MaTAR25* and confirm its function in human breast cancer cells. Based on syntenic conservation between the human and mouse genomes, we previously found three lincRNAs as potential human counterparts of *MaTAR25*: *LINC01270*, *LINC01271*, and *LINC01272* (29) (Fig. 6A). Among these three lncRNAs, only *LINC01271* is transcribed in the same direction as *MaTAR25*. Analysis of The Cancer Genome Atlas (TCGA) data (29) suggests two of these potential orthologs, *LINC01270* and *LINC01271*, are expressed at increased levels in multiple sub-types of breast cancer (Fig. 6A). Therefore, we focused on these two lincRNAs and performed independent ectopic expression of *LINC01271* and *LINC01270* in 4T1 *MaTAR25* KO cells to determine if one of these human lincRNAs could rescue the mouse *MaTAR25* KO phenotype. Cell viability assays indicated that ectopic expression of *LINC01271,* but not *LINC01270,* can rescue the proliferation phenotype of *MaTAR25* KO cells (Fig. 6B).

**Figure 6.**
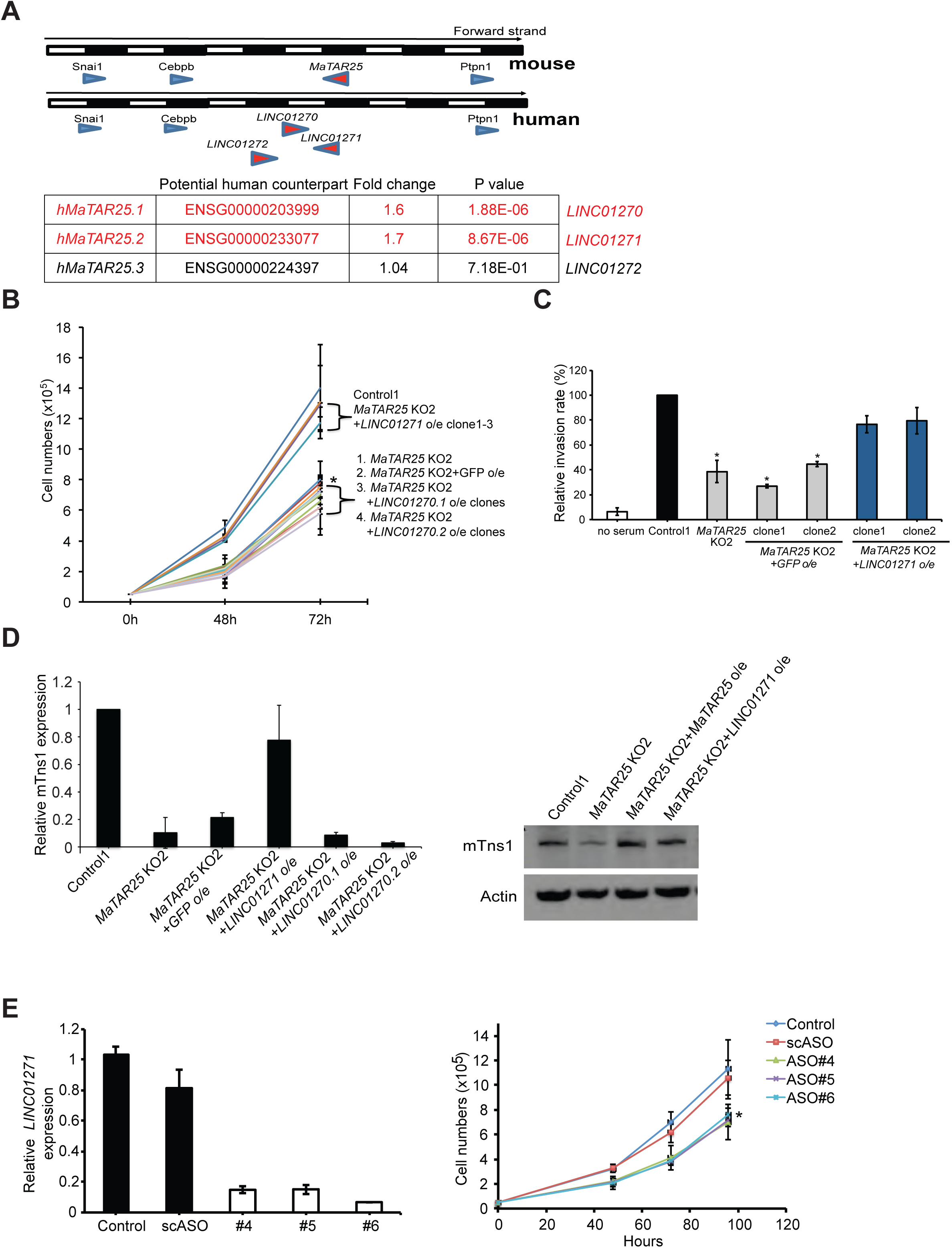
*LINC01271* is the human ortholog of *MaTAR25*. (A) All potential human orthologs of *MaTAR25* (*hMaTAR25*) were identified based on conservation of genomic location (synteny). RNA-seq data from The Cancer Genome Atlas (TCGA) was analyzed to evaluate the expression status of all potential *hMaTAR25* candidates by comparison of 1128 TCGA breast tumor datasets to 113 normal breast tissue controls. Fold change and statistical significance was calculated for the entire data matrix using DESeq2 (29). (B) Attempted rescue of 4T1 *MaTAR25* KO cells upon independent ectopic expression of two transcript isoforms of *LINC01270* (*LINC01270.1*, *LINC01270.2*), *or LINC01271* in cell viability assays. The mean cell numbers of three independent replicates of 4T1 control, *MaTAR25* KO, *MaTAR25* KO with GFP, *LINC01270.1*, *LINC01270.2*, and *LINC01271* is shown ± SD (n=3) \**p* < 0.05 (student’s t-test). Only ectopic expression of *LINC01271* can rescue the *MaTAR25* KO cell viability phenotype. (C) 4T1 *MaTAR25* KO cells with ectopic expression of GFP was used as a control to assess rescue in a cell invasion assay. The mean relative cell invasion of two independent replicates of 4T1 control, *MaTAR25* KO, *MaTAR25* KO with GFP, *LINC01271* ectopic expression is shown ± SD (n=2) \**p* < 0.05 (student’s t-test). Ectopic expression of *LINC01271* can rescue the *MaTAR25* KO cell invasion phenotype. (D) RNA expression level of *Tns1* was determined in *MaTAR25* KO cells ectopically expressing *LINC01270.1*, *LINC01270.2*, or *LINC01271* by qRT-PCR. The protein level of TNS1 was also examined in *MaTAR25* KO cells with ectopic expression of *LINC01271* by immunoblot analysis. (E) Three different ASOs targeting *LINC01271* were used to independently knockdown *LINC01271* in MDA-MB-231 LM2 cells. Left panel: the knockdown efficiency is shown ± SD (n=3) by qRT-PCR after 24 hours treatment of ASOs. Cells were seeded at the same density (5×10^4^/well) in 12-well tissue culture plates at day 0, ASOs were added to the culture medium and cell counting was performed at different time points to measure cell numbers. Right panel: The mean cell numbers of three independent replicates of MDA-MB-231 LM2 mock treated control cells, MDA-MB-231 LM2 cells treated with scrambled ASO, MDA-MB-231 LM2 cells independently treated with 3 different *LINC01271* ASOs is shown ± SD (n=3) \**p* < 0.05 (student’s t-test).

Invasion assays also showed that ectopic expression of *LINC01271* in 4T1 *MaTAR25* KO cells can rescue the cell invasion phenotype (Fig. 6C). In addition, the expression of *Tns1* can also be restored to a similar level as in 4T1 control cells upon overexpression of *LINC01271* in *MaTAR25* KO cells (Fig. 6D). Immunoprecipitation (RIP) using the PURB antibody indicated a specific interaction between PURB and *LINC01271* (Supplementary Fig. S6A) in human triple negative breast cancer MDA-MB-231 LM2 cells (44). Next, we performed smRNA-FISH to examine the localization of *LINC01271* within MDA-MB-231 LM2 cells and we found that similar to *MaTAR25* it is a nuclear enriched RNA (Supplementary Fig.S6C). Together, these results validate *LINC01271* as the human ortholog of *MaTAR25*.

### *LINC01271* may play a role in human breast cancer progression and have diagnostic and/or therapeutic potential

We performed ASO mediated KD of *LINC01271* in MDA-MB-231 LM2 cells and selected the three most effective *LINC01271* ASOs to assess a KD phenotype. After 96 hours of ASO treatment to mediate KD of *LINC01271* in MDA-MB-231 LM2 cells, all three independent ASOs decreased cell viability by approximately 32% (Fig. 6E). The KD result supports a role for *LINC01271* in human breast cancer progression.

According to the lncRNA database TANRIC (45), higher expression of *LINC01271* is correlated with poor breast cancer patient survival (Supplementary Fig. S6D). qRT-PCR analysis of the expression level of *LINC01271* in breast tumor organoids vs organoids derived from normal adjacent breast tissue showed a higher expression level of *LINC01271* in tumor-derived organoids (Supplementary Fig. S6E).

Next, we performed smRNA-FISH to localize *LINC01271* in patient breast tumor sections. We found that *LINC01271* expression level was increased with increased breast tumor stage (Fig. 7A and Supplementary Fig. S7A), and we identified the presence of clonal and regional differential expression patterns in most of the breast tumor patient samples (Supplementary Fig. S7B). Most interestingly, lung metastases exhibited higher expression of *LINC01271* than primary tumors from the same patients (Fig. 7B and Supplementary Fig. S7C). Thus, together these findings from patient-derived samples support our hypothesis that *LINC01271* is a potential therapeutic target to impact breast cancer progression and metastasis.

**Figure 7.**
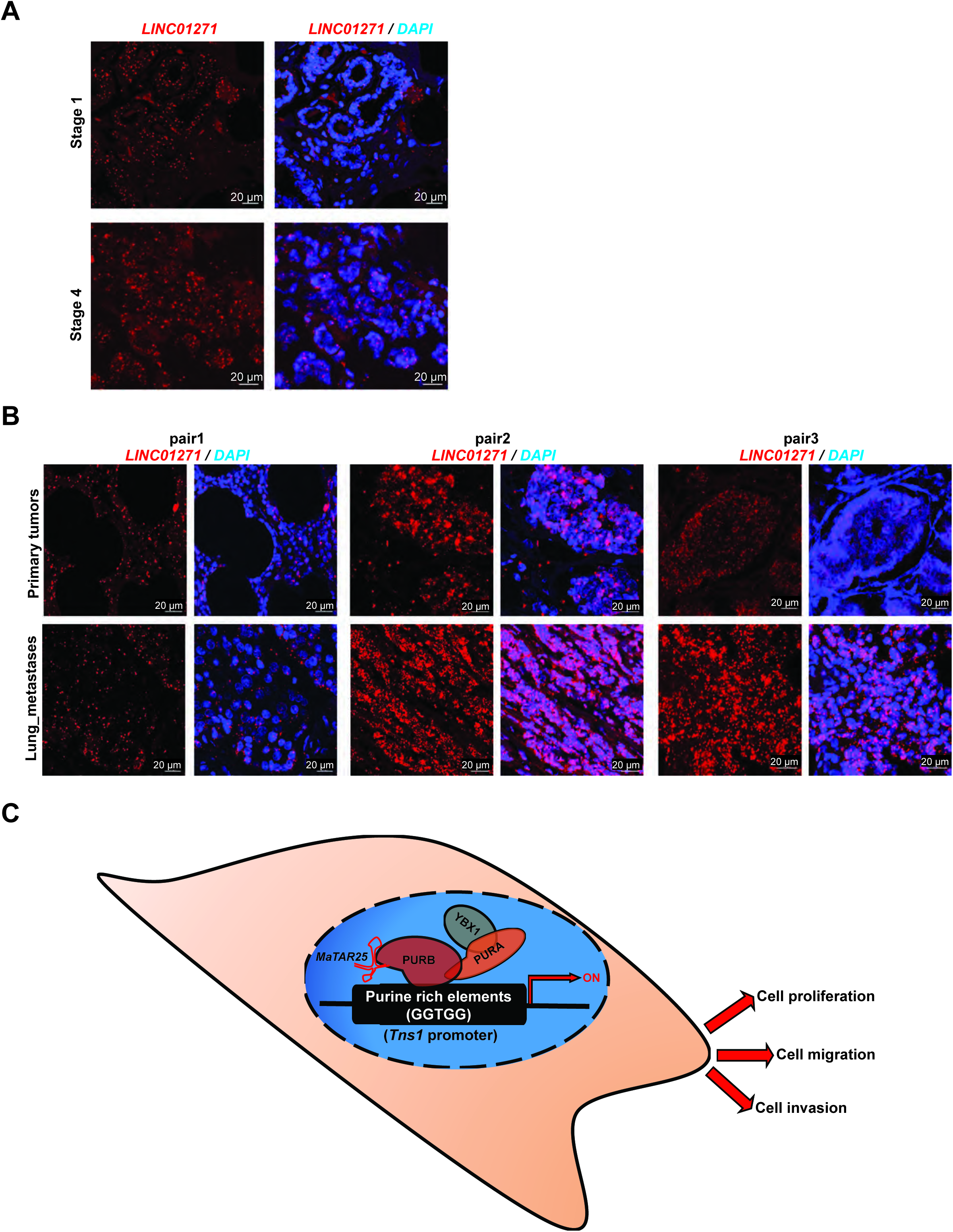
*LINC01271* expression in breast tumors and lung metastases. (A) Representative smRNA-FISH images showing the expression of *LINC01271* in human breast tumor sections from different stages of breast cancer. Scale bars are 20 μm. (B) Representative smRNA-FISH images showing the expression pattern of *LINC01271* within luminal subtype human primary breast tumors and lung metastases sections from the same patients. Scale bars are 20 μm. (C) Working model of *MaTAR25* function.

## DISCUSSION

We identified *MaTAR25* as a lncRNA that is upregulated in Her2+, luminal, and TNBC as compared to normal mammary epithelial cells. Genetic KO or ASO KD of *MaTAR25* results in a reduction in cell proliferation, migration, and invasion. Introduction of *MaTAR25* 4T1 KO cells into mice by mammary fat pad injection or tail vein injection results in smaller tumor growth and a significant reduction in lung metastases. The human ortholog of *MaTAR25* was identified as *LINC01271* and it is expressed in primary breast tumors and at even higher levels in lung metastases. Increased expression of the human ortholog of *MaTAR25* is associated with poor patient prognosis (44).

### *MaTAR25* regulates expression of the *Tns1* gene

In order to elucidate the molecular mechanism by which *MaTAR25* imparts its function, we used ChIRP-seq and RNA-seq to identify and validate the *Tns1* gene as a direct downstream target of *MaTAR25*. *MaTAR25* positively regulates the expression of *Tns1* in mammary tumor cells through binding to its DNA sequence in *trans*. Tns1 has been shown to localize to focal adhesions and to assist in mediating signaling between the extracellular matrix and the actin cytoskeleton to impact cell movement and proliferation (35, 36). Several reports have shown that loss of *Tns1* can cause a decrease in cell motility in many cell types (46–48), and *Tns1* has also been shown to be involved in epithelial-mesenchymal transition (EMT) of cancer cells (49). The general relationship between the expression of *Tns1* and different stages/subtypes of breast cancer has been unclear. However, Kaplan-Meier survival analysis indicates that the expression of *Tns1* is increased in grade 3 breast tumors (37) supporting our findings of a positive role of *Tns1* in breast tumor progression mediated by *MaTAR25*. Although *Tns1* is the top target of *MaTAR25* with the highest statistical significance we cannot rule out the possibility that *MaTAR25* also regulates additional genes given the other candidates identified in our ChIRP-seq analysis. These candidates will be the focus of future studies.

### *MaTAR25* partners with purine rich element binding protein B

By performing RNA antisense oligonucleotide pull-downs (RAP) in 4T1 cells combined with iTRAQ mass spectrometry, to determine the absolute quantitation of *MaTAR25* associated proteins, we identified a specific interaction between *MaTAR25* and purine rich element binding protein B (PURB) which appears to be crucial for the downstream regulation of *MaTAR25*. PURB has been reported to be a transcriptional co-activator and binds to the single strand of the repeated purine-rich element PUR which is present in promoter regions and as such has been implicated in transcriptional control. PURB plays different roles in many physiological and pathological processes (50–52). For example, PURB has been shown to be over-expressed in several different cancer types (53). In addition, a previous study showed PURB to act as a transcriptional co-factor that can be recruited by *linc-HOXA1* to mediate its transcriptional regulation in embryonic stem cells (43) supporting a critical role of PURB with different lincRNAs in different biological contexts. We identified the *MaTAR25* target region of the *Tns1* gene to have high ratio of purine and pyrimidine bases (3:1), and our ChIP-qPCR result indicated a high occupancy capacity of PURB over this targeting region, further supporting a functional role of PURB in partnering with *MaTAR25* to regulate the *Tns1* gene. The results of ectopic over-expression or siRNA mediated knockdown of PURB, resulting in altered expression (upregulation or down-regulation, respectively) of *Tns1* in 4T1 mammary tumor cells indicates a transcriptional regulatory role of PURB in this context. As the interaction of PURB with *MaTAR25* is essential for PURB binding to *Tns1* DNA this suggests that *MaTAR25* acts as a chaperone and/or scaffold for the *MaTAR25*/PURB/*Tns1* DNA complex, which is critical for transcriptional regulation of *Tns1* thereby impacting cancer progression.

PURB can form a homodimer, a heterodimer with PURA, or a heterotrimer with PURA and Y-box protein 1 (YBX1). Interestingly, these two additional proteins were also identified in our MS-iTRAQ analysis of *MaTAR25* interactors, and have been studied in many cancer types (54–56). Future investigation of these potential interactions may provide further insights into the molecular mechanism of the *MaTAR25* PURB complex in cancer cells.

### *LINC01271* is the human ortholog of *MaTAR25*

Based upon synteny and further validation we identified *LINC01271* as the human ortholog of *MaTAR25.* Interestingly, *LINC01271* has been identified as one of 65 new genetic loci that are related to overall breast cancer risk (57). *LINC01271* expression is increased in breast cancer and correlates with poor clinical outcome based on the analysis of patient clinical data (44). In addition, our results from examining sections of breast tumors and corresponding lung metastasis from the same patients showed a positive correlation between the high expression of *LINC01271* and breast cancer stage. Our finding of an even higher level of expression in lung metastases from the same patients was especially interesting. Metastasis is the major cause of cancer related deaths, particularly in breast cancer patients (58, 59), and developing an efficient treatment strategy to target and reduce breast cancer metastasis still remains the key challenge for this disease. Our ability to use ASOs to knockdown *MaTAR25* in the Her2/neu mouse model resulting in necrotic tumors and a significant reduction in metastasis has therapeutic implications. ASO targeting as a therapeutic approach has been applied to many diseases (60–62) including cancer (26, 63, 64) in recent years. For example, a cEt ASO targeting the transcription factor STAT3 has shown robust single agent activity in highly treatment-refractory lymphoma and non-small cell lung cancer studies (64). The STAT3 ASO (AZD9150) has advanced into multiple Phase I and II clinical trials (NCT01563302, NCT02983578, NCT02549651). In addition, an antisense drug targeting all forms of the androgen receptor for the treatment of advanced metastatic prostate cancer has entered a clinical trial (NCT02144051). In this study, we developed three ASOs targeting *LINC01271* for functional assays *in vitro*. Ultimately, *LINC01271* represents an ideal candidate to be exploited as a potential prognostic/therapeutic target and *LINC01271* specific ASOs as therapeutics to impact breast cancer progression and metastasis.

## METHODS EXPERIMENTAL MODELS

### Cell Culture

Murine 4T1 cells, murine NF639 (MMTV-cNEU) cells, human MDA-MB-231 cells, and human MDA-MB-231 LM2 cells were cultured in DMEM with 10% FBS, and 1% penicillin/strptomycin.

### Organoid Culture

Surgically removed tumor samples from breast cancer patients along with adjacent normal tissue were collected from Northwell Health in accordance with Institutional Review Board protocol IRB-03-012 (TAP16-08). Informed consent ensured that the de-identified materials collected, the models created, and data generated from them can be shared without exceptions with researchers in the scientific community. Tumor and normal organoids were developed using a previously published protocol (Sachs et al., 2018). The tissues were manually cut into smaller pieces and treated with Collagenase IV at 37°C. The samples were manually broken down by pipetting into smaller fragments and seeded in a dome of matrigel. Organoids were grown in culture media which contained 10% R-Spondin 1 conditioned media, 5 nM Neuregulin 1, 5 ng/ml FGF7, 20 ng/ml FGF10, 5 ng/ml EGF, 100 ng/ml Noggin, 500 nM A83-01, 5 μM Y-27632, 1.2 μM SB 202190, 1x B27 supplement, 1.25 mM N-Acetyl-cysteine, 5 mM Nicotinamide, 1x Glutamax, 10 mM Hepes, 100U/ml Penicillin/streptomycin, 50 μg/ml Primocin in 1x Advanced DMEM-F12 (Sachs et al., 2018). Cultures were passaged every 2-4 weeks using TrypLETM to break down the organoids into smaller clusters of cells and replating them in Matrigel domes.

### Mice

All animal procedures and studies were approved by the Cold Spring Harbor Laboratory Animal Use Committee in accordance to IACUC procedures. Briefly, male MMTV-Neu-NDL mice (FVB/N background) were kindly provided by Dr. William Muller (McGill University, Canada). Male MMTV-Neu-NDL mice were crossed with wild type FVB/N female mice purchased from the Jackson Laboratory for breeding. PCR genotyping was applied to select female mice with heterogeneous genotypes for MMTV-Neu-NDL for later *in vivo* ASO injection experiments. 4-6 week old BALB/c female mice were purchased from the Jackson Laboratory for *in vivo* 4T1 cell mammary fat pad (MFP) injection and tail-vein injection experiments.

## METHODS

Details of materials and reagents are listed in Supplementary Table S2

Details of oligonucleotide sequences are listed in Supplementary Table S3

### 5’/3’ Rapid Amplification of cDNA Ends (RACE)

5’ and 3’ RACE of *MaTAR25* transcripts was performed on TRIzol-extracted RNA using the Ambion FirstChoice RLM-RACE kit according to the manufacturer’s instructions. Briefly, fragments were amplified by nested PCR using AmpliTaq Polymerase. PCR products were separated on 2% agarose, bands excised, gel purified, sub-cloned into pGEM-T Easy (Promega) and 4 or more clones per fragment were sequenced using standard Sanger sequencing. Primer sequences are provided in Table S1.

### Northern Blot Analysis

*MaTAR25*-specific radiolabeled DNA probes were generated using dCTP P32 in a random primed labeling reaction. Total RNA was extracted by the TRIzol method. Analysis of RNA expression was performed by following NorthernMax® Kit manual. Briefly, 20 μg and 30 μg total RNA samples were electrophoresed on a 1% agarose gel and was transferred to a positively charged nylon membrane (NC). The RNA was then fixed to the NC membrane using UV crosslinking. The cross-linked membrane was prehybridized with ultrahyb-oligo hybridization buffer and hybridized with the *MaTAR25*-specific radiolabeled DNA probe. After washing with SSC wash buffers several times, the wrapped membrane was exposed to a PhosphorImager screen in a cassette or X ray film for detecting signals.

### Cell Lysate Preparation for Immunoblot Analysis

Cells were trypsinized, and harvested cell pellets were lysed in RIPA buffer (25 mM Tris-HCl pH 7.6, 150 mM NaCl, 1% NP-40 substitute, 1% sodium deoxycholate and 0.1% SDS) supplemented with 1X Roche protease inhibitor cocktail. The cell lysate was incubated on ice for 15 minutes, then sonicated for 5 minutes before centrifugation at 13000xg. The supernatant was collected and quantified using the BCA protein assay.

### *In vitro* Transcription/Translation

T7 promoter containing DNA or plasmids were used in a TNT® Quick Coupled Transcription/Translation System following the manufacture’s protocol. Briefly, 1 μg Plasmid DNA Template was mixed with 40 μl TNT® T7 Quick Master Mix, 1 μl Methionine (1 mM), 1 μl Transcend™ Biotin-Lysyl-tRNA, and nuclease free water for the final volume of 50 μl per reaction. The reaction tube was incubated at 30°C for 90 minutes, and 1 μl reaction product was added into diluted 2x Laemmli sample buffer for immunoblot analysis. Samples were loaded on 4–20% Mini-PROTEAN® TGX™ Precast Protein Gels, and the signals were detected by Streptavidin-HRP.

### RNA Isolation and Quantitative Real-Time PCR (qRT-PCR) Assays

Total RNA was extracted using TRIzol following the manufacture’s protocol. 1 μg total RNA was treated with DNaseI and reverse transcribed into cDNA using TaqMan Reverse Transcription Reagent kit, followed by qPCR with SYBR green PCR master mix on an ABI QuantStudio 6 Flex Real-Time PCR System. qRT-PCR conditions were as follows: 30 minutes at 50°C for reverse transcription, 15 minutes at 95°C for the initial activation step followed by 40 cycles of 15 seconds at 94°C, 30 seconds at 60°C.

Mouse peptidylprolyl isomerase B (cyclophilin B; PPIB), and human GAPDH and RPL13A were used as endogenous controls to normalize each sample. A list of primers used is provided in Table S1.

### Cell Fractionation, Cytoplasmic/Nucleoplasmic/Chromatin-related RNA Isolation

Cell fractionation was done using a standardized protocol previously described (66). Briefly, cultured cells were harvested and lysed in NP-40 substitute lysis buffer (10 mM Tris pH 7.5, 150 mM NaCl, and 0.15% NP-40 substitute). The cell lysate was overlaid on top of sucrose buffer (10 mM Tris pH7.4, 150 mM NaCl, and 24% sucrose) and centrifuged at 3500xg for 10 minutes to separate the cytoplasmic fraction and nuclei pellet. The nuclei pellet was rinsed with PBS-EDTA once, and resuspended in glycerol buffer (20 mM Tris pH7.4, 75 mM NaCl, 0.5 mM EDTA, and 50% Glycerol) mixed with Urea buffer (1 M Urea, 0.3 M NaCl, 7.5 mM MgCl_2_, 0.2 mM EDTA, and 1% NP-40 substitute) on ice for 2 minutes. The lysate was centrifuged at 13000xg for 2 minutes to separate the nucleoplasmic fraction and chromatin pellet. The chromatin later was resuspended in TRIzol reagent and fully solubilized by passing through the 21-gauge needle and syringe. The cytoplasmic fraction and nucleoplasmic fraction were also used for RNA extraction using TRIzol reagent. RNA extracted from different fractions were applied for cDNA synthesis and qRT-PCR. Primer sequences are provided in Table S1.

### CRISPR/Cas9 Genetic Knockout

To generate a genetic knockout of *MaTAR25*, two sgRNAs targeting the promoter region were combined, creating a deletion including the TSS. Both sgRNAs were designed using http://crispr.mit.edu/. The sgRNA targeting the gene body of *MaTAR25* was cloned into a pSpCas9(BB)-2A-GFP vector and the sgRNA targeting the upstream promoter region was cloned into a pSpCas9(BB)-2A-mCherry vector. 4T1 cells were transfected with both plasmids using 4D-Nucleofector X Unit, using program code “CN-114”. To select for cells expressing both gRNAs, GFP and mCherry double positive cells were sorted 40 hours post transfection, as single cell deposition into 96-well plates using a FACS Aria (SORP) Cell Sorter (BD). Each single cell clone was propagated and analyzed by genomic PCR and qRT-PCR to select for homozygous knockout clones. Cells transfected with a sgRNA targeting *Renilla* luciferase were used as a negative control. Sequences for sgRNAs and primers are provided in Table S1.

### Cell Counting Viability Assay

Cultured cells were harvested and the same number of cells were seeded into each well of a 12 well tissue culture plate at day 0. Trypan Blue-treated cell suspensions were collected and applied to a hemocytometer for manual counting at different time points.

### Cell Cycle Analysis

Cell cycle analysis was performed using BD bromodeoxyuridine (BrdU) FITC assay kit following the manufacturer’s protocol. Briefly, cultured cells were incubated with BrdU containing medium for 30 minutes, and FITC conjugated anti-BrdU antibody was applied for labeling actively synthesizing DNA. 7-aminoactinomycin D (7-AAD) was used for labeling total DNA. The labeled cell samples were analyzed on the BD LSR II flow cytometer, and BrdU FITC-A vs DNA 7-AAD dot plot with gates was used to encompass the G0/G1, S, and G2/M populations. The collected cytometry data were analyzed with FACSDiva™ and FlowJo software.

### Migration Assay

Live cell tracking was performed to examine cell migration. Cultured cells were harvested and the same number of cells were seeded into 6 wells of a tissue culture plate at day 0. Images were collected every 5 minutes (5 viewpoints were selected from each well) using Zeiss AxioObserver microscope for 8 hours, and the images of individual sample were converted to videos using ImageJ. Videos were analyzed by CellTracker image processing software. The mean relative migration distance (μm) of three independent replicates of 4T1 control groups and *MaTAR25* KO groups were calculated.

### Scratch Wound Healing Assay

Cultured cells were harvested and seeded into each well of a 24 well tissue culture plate. Cells were incubated until they reached at least 90% confluence. The wound line was created by “scratch” with a p200 micropipette tip, then cells were washed with PBS twice to remove the debris and then each well was imaged. After 12 hours incubation, each well was images again and the migration areas in each well was measured by ImageJ for comparison.

### Invasion Assay

Invasion assays with 4T1 *MaTAR25* knockout cells and 4T1 control cells were carried out using the Cultrex® 24 well BME Cell Invasion Assay (Trevigen) following the manufacturer’s protocol. Briefly, cells were harvested and seeded at a density of 1 x 10^5^ cells/well into the invasion chamber. As a negative control, serum-free medium was used that did not stimulate cell invasion through the BME. The plate was incubated at 37°C for 24 hours and the assay was performed. The tumor cells that invaded through the BME layer and attached to the bottom of the invasion chamber were collected using cell dissociation solution and stained with Calcein AM solution. The fluorescence was measured with a SpectraMax i3 Multi-Mode Detection Platform (Molecular Devices) using the 480/520 nm filter set.

### Cloning

Specific gene overexpression plasmids were constructed using NEBuilder HiFi DNA Assembly following the manufacture’s protocol. PCR products were cloned into pCMV6-entry plasmids digested with Sgf I and Fse I. Assembled plasmids were introduced into NEB stable competent E. *coli* using heat shock transformation and kanamycin selection.

4 or more colonies per plate were picked and sequenced using standard Sanger sequencing.

### Antisense Oligonucleotide (ASO) and siRNA-Mediated Knockdown (KD)

Specific 16mer antisense oligonucleotides (ASOs) comprised of phosphorothioate-modified short S-cEt (S-2 ′ -O-Et-2 ′, 4′ -bridged nucleic acid) gapmer chemistry targeting *MaTAR25* and *LINC01271* were designed and provided by Ionis Pharmaceuticals, Inc. Briefly, cultured cells were harvested and seeded into culture dishes. Transfection-free uptake of ASOs was accomplished by adding 4 μM of either *MaTAR25/LINC01271*-specific ASOs or scrambled ASO (scASO) to the culture medium immediately after seeding the cells. Cells were incubated for indicated time points and RNA was isolated using TRIzol reagent for qRT-PCR to check the knockdown efficiency. For siRNA mediated knockdown, siRNAs (27mers) targeting mouse PURB were purchased from ORIGENE, and siRNA transfection was done using Lipofectamine 2000 following the manufacture’s protocol. RNA was extracted at different time points for qRT-PCR to check the knockdown efficiency. ASO sequences and primer sequences are provided in Table S1.

### Chromatin Isolation by RNA purification (ChIRP)-Seq

For Chromatin Isolation by RNA purification (ChIRP), we followed a previously described protocol (34). Briefly, 20 million cells were harvested and fixed in 1% glutaraldehyde solution for each reaction. ChIRP was performed using biotinylated oligo probes designed against mouse *MaTAR25* using the ChIRP probe designer (Biosearch Technologies). Independent even and odd probe pools were used to ensure lncRNA-specific retrieval (refer to Table S1 for odd and even sequences targeting *MaTAR25*, and probes targeting mouse PPIB transcripts which were used as negative controls). ChIRP-Seq libraries were constructed using the Illumina TruSeq ChIP Library Preparation Kit. Sequencing libraries were barcoded using TruSeq adapters and sequenced on Illumina NextSeq instruments.

### ChIRP-Seq Data analysis

Data quality was assessed using FastQC (http://www.bioinformatics.babraham.ac.uk/projects/fastqc/) and paired-end reads were mapped to GRCm38 using Bowtie2 (67) with parameters --end-to-end --sensitive --fr, resulting in a 90% or higher overall alignment rate. ChIRP seq analysis was performed using HOMER (65). Differential ChIRP peaks were called using the getDifferentialPeaksReplicates.pl script, with negative control (PPIB) pull-down samples as background and parameters -style histone -f 50. Peaks identified with at least a 50-fold enrichment were processed further using the annotatePeaks.pl script and the GENCODE vM16 annotation. Both known and *de novo* motif analysis was carried out with the findMotifsGenome.pl script on the repeat-masked GRCm38 genome, +/-500 bp around the identified ChIRP peaks.

### RNA-seq Library Construction

RNA was extracted using TRIzol following the manufacture’s protocol. RNA quality was assayed by running an RNA 6000 Nano chip on a 2100 Bioanalyzer. 1 μg total RNA was used for constructing each RNA-seq library using the Illumina TruSeq sample prep kit v2 following the manufacture’s protocol. Briefly, RNA was polyA selected and enzymatically fragmented. cDNA was synthesized using Super Script II master mix, followed by end repair, A-tailing and PCR amplification. Each library was high-throughput single-end sequenced on Illumina NextSeq instruments.

### RNA-Seq Data analysis

Data was analyzed as previously described (29). Briefly, the quality of FASTQ files was assessed using FastQC (http://www.bioinformatics.babraham.ac.uk/projects/fastqc/). Reads were mapped to GRCm38 using STAR (68), and the reads per gene record were counted using HTSeq-count (69) and the GENCODE vM5 annotation. Differential gene expression was performed with DESeq2 (70), and an adjusted p-value of < 0.05 was set as threshold for statistical significance. KEGG pathway and GO term enrichment and was carried out using the R/Bioconductor packages GAGE 71) and Pathview (72).

### RNA Antisense Pulldown and Mass Spectrometry

Cells were lysed in a 10 cm culture dish in 1 ml IP lysis buffer (IPLB, 25 mM Tris-HCl pH 7.4, 150 mM NaCl, 1% NP-40, 1 mM EDTA, 5% glycerol, supplemented with 100 U/ml SUPERase-IN and 1X Roche protease inhibitor cocktail) for 10 minutes, and lysate was centrifuged at 13,000xg for 10 minutes. Cell lysate was adjusted to 0.3 mg/ml (Pierce BCA Protein Assay). A total of 100 pmol of biotinylated oligo was added to 500 μl of lysate and incubated at room temperature for 1 hour with rotation. 100 μl streptavidin Dynabeads were washed in IPLB, added to the lysate, and incubated for 30 minutes at room temperature with rotation. Beads were washed three times with 1 ml lysis buffer. For determining temperature for optimal elution, beads were then resuspended in 240 μl of 100 mM TEAB and aliquoted into eight PCR tubes. Temperature was set on a veriflex PCR block and incubated for 10 minutes. Beads were captured and TRIzol was added to the eluate and beads. Once optimal temperature is established, the beads were resuspended in 90 μl of 100 mM TEAB, and incubated at 50° C for 10 minutes. TRIzol was added to 30 μl of the eluate, another 30 μl was kept for immunoblots, and the last 30 μl aliquot was sent directly to the Cold Spring Harbor Laboratory Mass Spectrometry Shared Resource for analysis.

### RNA Immunoprecipitation (RIP)

RIP was performed following RIP the Abcam protocol with minor modifications. Briefly, cultured cells were harvested and 40 million cells were washed once with cold PBS, then the cells were resuspended in 8 ml PBS, 8 ml nuclear isolation buffer (1.28 M sucrose, 40 mM Tris-HCl pH 7.5, 20 mM MgCl_2_, and 4% Triton X-100 supplemented with 100 U/ml SUPERase-IN and 1X Roche protease inhibitor cocktail), and 24 ml nuclease free water on ice for 20 min with frequent mixing. The cleared lysates were pelleted by centrifugation at 2,500xg for 15 min. Pellets resuspended in 4 ml RIP buffer (150 mM KCl, 25 mM Tris pH 7.4, 5 mM EDTA, 0.5 mM DTT, and 0.5% NP40 substitute supplemented with 100 U/ml SUPERase-IN and 1X Roche protease inhibitor cocktail) and sonicated for 5 minutes using BioRuptor Pico water bath sonicator (30 s ON/OFF) at 4°C. The lysates were cleaned by centrifugation at 13,000 rpm for 10 minutes. The supertanant was collected and separated, then incubated with 4 μg PURB antibody or rabbit isotype IgG control for 2 hours to overnight at 4°C with gentle rotation. 80 μl of protein A beads for rabbit antibody then added into the reactions and incubate for 1 hour at 4°C with gentle rotation. After washing three times with RIP buffer and once with

PBS, beads were collected for immunoblot analysis and RNA extraction for qRT-PCR. Primers for RIP qRT-PCR can be found in Table S1.

### Chromatin Immunoprecipitation (ChIP) coupled with quantitative PCR (ChIP-qPCR)

For Chromatin Immunoprecipitation (ChIP) we followed protocols previously described (73). Briefly, 30 million 4T1 cells were harvested and crosslinked in 1% formaldehyde at room temperature for 20 minutes, then the reaction was quenched using 0.125 M glycine. Cells were incubated with cell lysis buffer (10 mM Tris pH8.0, 10 mM NaCl, 0.2% NP-40 substitute) and then resuspended and sonicated in 1.5 ml of nuclei lysis buffer (50 mM Tris pH8.0, 10 mM EDTA, 1% SDS) for 15 min using BioRuptor Pico water bath sonicator (30 s ON/OFF) at 4°C. For one IP, 1.5 ml of sonicated chromatin from 30 million cells were diluted with 21 ml IP-Dilution buffer (20 mM Tris pH 8.0, 2 mM EDTA, 150 mM NaCl, 1% Triton X-100, 0.01% SDS,) and incubated with 4 μg of PURB antibody or rabbit isotype IgG control, and 80 *μ*l of protein A beads for rabbit antibody at 4°C overnight. After washing once with IP-wash 1 buffer (20 mM Tris pH8.0, 2 mM EDTA, 50 mM NaCl, 1% Triton X-100, 0.1% SDS), twice with High-salt buffer (20 mM Tris pH 8.0, 2 mM EDTA, 500 mM NaCl, 1% Triton X-100, 0.01% SDS), once with IP-wash 2 buffer (10 mM Tris pH 8.0, 1 mM EDTA 0.25 M LiCl, 1% NP-40 substitute, 1% sodium deoxycholate), twice with TE buffer (10 mM Tris-Cl, 1 mM EDTA, pH 8.0), beads bound chromatin were eluted in 800 μl nuclei lysis buffer by heating at 65 °C for 15 minutes. 48 μl of 5 M NaCl was added to the 800 *μ*l eluted chromatin, followed by incubation at 65°C overnight for reverse cross-linking. After reverse cross-linking, DNA was treated with RNaseA and proteinase K, followed by purification using QIAGEN PCR purification kit. qPCR was performed on ABI QuantStudio 6 Flex Real-Time PCR System. ChIP-qPCR primers can be found in Table S1.

### Single Molecule RNA Fluorescence In Situ Hybridization (FISH)

For single-molecule RNA FISH, custom Type-6 primary probes targeting *MaTAR25*, *LINC01271* and other lncRNAs were designed and synthesized by Affymetrix. For RNA-FISH on cultured cell samples, Affymetrix View ISH Cell Assay Kit reagents were used. Cultured cells were harvested and seeded onto acid-cleaned #1.5 glass coverslips for 24 hours incubation to 70% confluence, cell samples then were fixed in freshly-prepared 4% paraformaldehyde (PFA). Cells were then permeabilized and protease digested before hybridization. For RNA-FISH on formalin-fixed paraffin-embedded (FFPE) tissue sections of breast tumors and metastases, Affymetrix ViewRNA ISH Tissue 1-Plex Assay kit reagents were applied. Sections on slides were deparaffinized, protease digested, and fixed with 10% NBF before hybridization. QuantiGene ViewRNA probe hybridizations were performed at 40°C for 3 hours. The hybridization and signal amplification steps were performed according to the manufacturer’s instructions, and nuclei were counter-stained with DAPI. Coverslips and tissue sections were mounted in ProLong Gold Antifade mounting medium before detection. Imaging was performed on Zeiss LSM 710/780 Confocal Microscope systems.

### DNA FISH

Different mouse BAC clones (RPCI-23) were used as template included (*MaTAR25*), and (*Tensin1*). 1 μg BAC DNA was used as template for random priming reaction to generate amine-modified DNA, and amine-modified DNA was labelled with a reactive fluorescent dye as fluorescent probes according to the protocol provided with ARES™ Alexa Fluor™ DNA Labeling Kit. For DNA FISH, we followed protocols previously described (Hogan et al., 2015). Briefly, cultured cells were seeded onto 22mm^2^ glass coverslips (Corning), and coverslips were fixed with freshly prepared 4% PFA for 20 minutes at room temperature, and permeablized in 0.5% Triton X-100/1X PBS for 5 minutes on ice. Probes were prepared for hybridization by mixing 2μl probe with 5μl each of sheared salmon sperm DNA, mouse Cot1 DNA, and yeast tRNA, dehydrating the probe mixture in the speed-vac, and then resuspending the probe in 10μl deionized formamide (Ambion). Just prior to hybridization, the probes were denatured at 95°C for 10 minutes, transferred to ice for 5 minutes, and then mixed with 10μl 2X Hybridization Buffer (4X SSC, 20% dextran sulfate) and pipetted onto slides so that coverslips could be placed cell-side-down on the probe mixture for the hybridization reaction. After several washes, nuclei were counter-stained with DAPI. Coverslips were mounted in ProLong Gold Antifade mounting medium before detection. Coverslips were imaged using Zeiss LSM 710/780 Confocal Microscope systems.

### Immunofluorescence (IF)

For IF, we followed protocols previously described with minor modifications depended on applied antibody (74). Briefly, cultured cells were harvested and seeded onto acid-cleaned #1.5 glass coverslips for 24 hours incubation to 70% confluence, cell samples then were fixed in 4% formaldehyde for 20 minutes. Samples were permeabilized in 0.2% Triton X-100 plus 1% Bovine Serum Albumin (BSA) in PBS for 5 minutes on ice. After incubated in 1% Bovine Serum Albumin (BSA) in PBS for 30 minutes blocking, samples were incubated in the appropriate concentration (1:50-1:200 followed by mamufacturer’s recommendations) of primary antibody for 1-2 hours at room temperature, and incubated in diluted secondary antibody solution (Alexa 594 conjugated) with phalloidin (Alexa 488 conjugated) for 1 hour in a humidified chamber at room temperature. After several washes, nuclei were counter-stained with DAPI. Coverslips were mounted in ProLong Gold Antifade mounting medium before detection. Coverslips were imaged on the Zeiss LSM 710/780 Confocal Microscope systems.

### Transmission Electron Microscopy (TEM)

Cultured cells were harvested and seeded onto 10 cm culture dishes for 24 hours, and fixed with 2.5% glutaraldehyde (EM grade) in 0.1 M phosphate buffer pH 7.4 at room temperature for 1 hour. The fixed cells were collected by using a cell scraper, washed several times in 0.1M phosphate buffer and post fixed in 1% Osmium tetroxide. Samples were dehydrated in ethanol (30%, 50%, 70%, 80%, 95%, and 100%), embedding in EMbed 812 Resin and polymerized. 70-90 nm sections were cut on a Reichert-Jung ultramicrotome using a diamond knife (DiATOME). Sections were collected on copper grids, stained with UranyLess and lead citrate and imaged with a Hitachi H-7000 TEM.

### *In Vivo* Mouse Model ASO Injection

Three month old MMTV-Neu-NDL mice were divided into three cohorts (7-12 mice each), and each mouse in the cohort received either scASO or *MaTAR25* specific ASO1 or ASO2 via subcutaneous injection 50mg/kg/day twice per week. The injections were carried out for a period of 7 weeks, upon which at least one tumor from most of the control mice reached 2 cm in size. During the course of treatment, tumors were measured twice per week. At the end of the treatment period the animals were euthanized and the primary tumors, lungs, livers, spleens, and intestines were collected. Collected lungs were fixed in 4% paraformaldehyde and incubated in 20-30% sucrose solution overnight, then frozen in OCT solution. The lung OCT blocks were cross-sectioned 2 mm apart, and the lung sections were embedded horizontally to obtain serial sections of the entire lung. Other tissues were cut into two pieces. One parts of each issue was snap frozen using liquid nitrogen for further RNA extraction. The remaining tissues were fixed in 4% paraformaldehyde and paraffin embedded, then formalin-fixed, paraffin embeeded (FFPE) tissue blocks were sectioned. All sections were stained with Hematoxlin and Eosin (H&E) following standard protocol, and slides were scanned and analyzed using an Aperio ImageScope pathology slide viewing system. All samples were processed and stained at the Cold Spring Harbor Laboratory Histology Shared Resource.

### *In Vivo* 4T1 Cells Injection

5-6 week old female BALB/c mice and 4T1 cells (control and *MaTAR25* KO cells) expressing luciferase were used for 4T1 mammary fat pad and tail-vein injection experiments. For mammary fat pad injection, 1×10^5^ 4T1 control or *MaTAR25* KO cells were injected orthotopically into the mammary fat pad of female BALB/c mice. Mice were monitored and primary tumors were measured every week. Mice were sacrificed and tumors were collected at day 28 to compare the tumor growth rate between 4T1 control groups and *MaTAR25* KO groups.

For tail-vein injection, female BALB/c mice were injected intravenously with 1×10^5^ 4T1control or *MaTAR25* KO cells in the tail vein. Mice were monitored every week and sacrificed at day 21. The mouse lungs were collected and imaged, and lung metastatic nodules were counted to compare the metastatic ability between 4T1 control groups and *MaTAR25* KO groups.

## QUANTIFICATION AND STATISTICAL ANALYSIS

Statistics tests were performed and analysed using Microsoft Excel and GraphPad Prism 7.0. p value was calculated by paired Student’s t-test, two-tailed. Significance was defined as p < 0.05.

## DATA AND SOFTWARE AVAILABILITY

The accession number for the RNA-seq and ChIRP-seq data reported in this study is GEO: GSE142169

## Authors’ Contributions

**Conception and design**: K.Chang and D.L. Spector

Analysis of the RNA-seq, ChIRP-seq, and TCGA human BC patient data**: S.D. Diermeier**

**Cell cycle analysis, and technical support of CRISPR mediated KO and RNA antisense pulldown**: A.T. Yu

**Tissue RNA-FISH**: L.D. Brine

Technical support (*in vivo* mouse model ASO injections and 4T1 mammary fat pad injections)**: S. Russo**

**Human organoid culture and organoid RNA extraction**: S. Bhatia.

**TEM sample preparation and imaging:** H. Alsudani

**Material support (i.e., human BC patient samples for organoid culture and FFPE slides)**: K. Kostroff, T. Bhuiya, E. Brogi

Material support (designing and providing ASOs for *in vitro* and *in vivo* **experiments)**: C. F. Bennett, F. Rigo

**Writing of the manuscript**: K.Chang and D.L. Spector

**Study supervision**: D.L. Spector

## Acknowledgments

We thank members of the Spector lab for critical discussions and advice throughout the course of this study. We also acknowledge Dr. J. Erby Wilkinson (University of Michigan Medical School) for help with pathology and histology analysis. We thank Dr. William Muller (McGill University, Montreal) for providing MMTV-Neu-NDL mice, Dr. Mikala Egeblad (CSHL) for providing MMTV-PyMT mice, Dr. Fred R. Miller (Wayne State University) for providing 4T1 cells, Dr. Nicholas Tonks for providing SKBR3 and BT-474 cells, Dr. Joan Massagué (Sloan Kettering Institute) for providing MDA-MB-231 LM2 cells, Dr. Bruce Stillman (CSHL) for providing the pET3a-Xenopus laevis histone H2B plasmid, and Dr. Scott Lyons (CSHL) for providing Ef1α-Luc2-mStrawberry plasmid. We would also like to thank the CSHL Cancer Center Shared Resources (Microscopy, Flow Cytometry, Animal, Histology, and Next-Gen Sequencing) for services and technical expertise (NCI 2P3OCA45508). This research was supported by NCI 5P01CA013106-Project 3 (D.L.S.), Susan G. Komen postdoctoral fellowship (S.D.D.) and NCI 1K99CA215362 (S.D.D.), CSHL/Northwell Health (D.L.S.), and Manhasset Woman’s Coalition Against Breast Cancer (S.B.).

