## Supplementary Figures for "*MaTAR25* LncRNA Regulates the *Tensin1* Gene to Impact Breast Cancer Progression"

Supp. Fig. 1

A

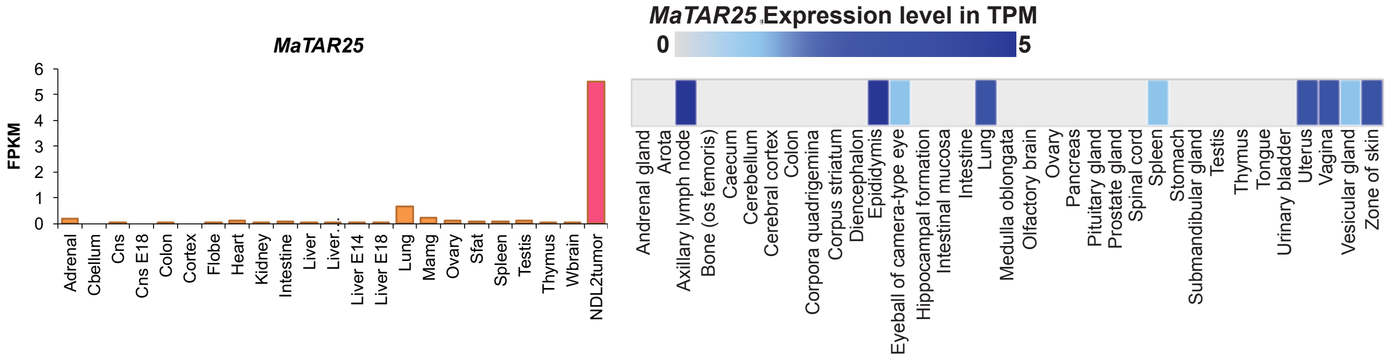


B

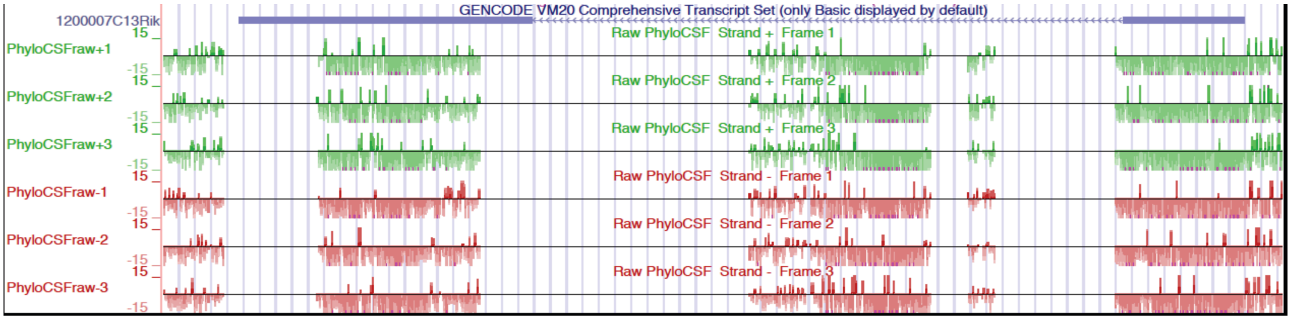


C

| Result for species name : mm10 with job ID :1563826412 | | | | | | | |
|---|---|---|---|---|---|---|---|
| Data ID | Sequence Name | RNA Size | ORF Size | Ficket Score | Hexamer Score | Coding Probability | Coding Label |
| 0 | MATAR25 | 1978 | 369 | 0.8146 | 0.00331186776961 | 0.24503635507318 | no |

D

| ID | C/NC | CODING POTENTIAL SCORE | EVIDENCE | UTR-DB HITs | RNA-DB HITs |
|---|---|---|---|---|---|
| MaTAR25 | noncoding (weak) | -0.932493 | detail | search | search |

**ORF INFORMATION**

*unreliable ORF*

| SOURCE | START | END | LENGTH | COVERAGE | SCORE | TYPE |
|---|---|---|---|---|---|---|
| ORF_FRAMEFINDER | 1121 | 1487 | 367 (123AA) | 18.50% | 46.34 | Full |

**PUTATIVE PEPTIDE**

*unreliable ORF*

>MaTAR25 [framefinder (1120,1486) score=46.34 used=18.50% {forward,strict} ]

```
MMTGDIPSLPWGEGGLWPGDKGSRPPTSGKRTDNRDSGQATSPGEGGHRMSVREDRDRGC
LMRPSARPTASLPGSRPLSCEVEPPFSLFYHPSSSQLAGVPSRCPLACGLGSEGKGTNPW
VN
```

E

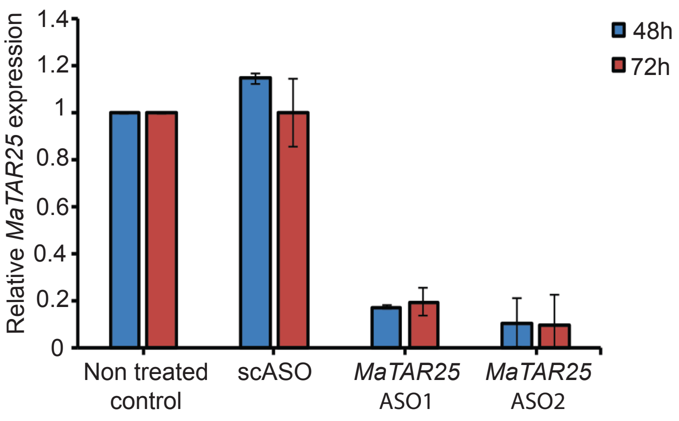


F

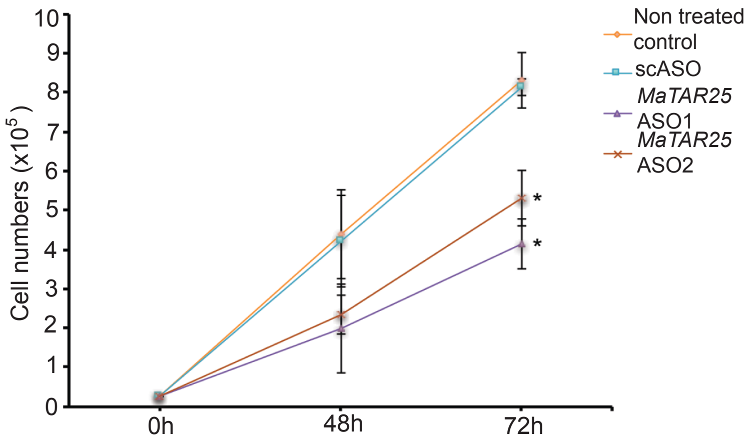

Supp. Fig. 2

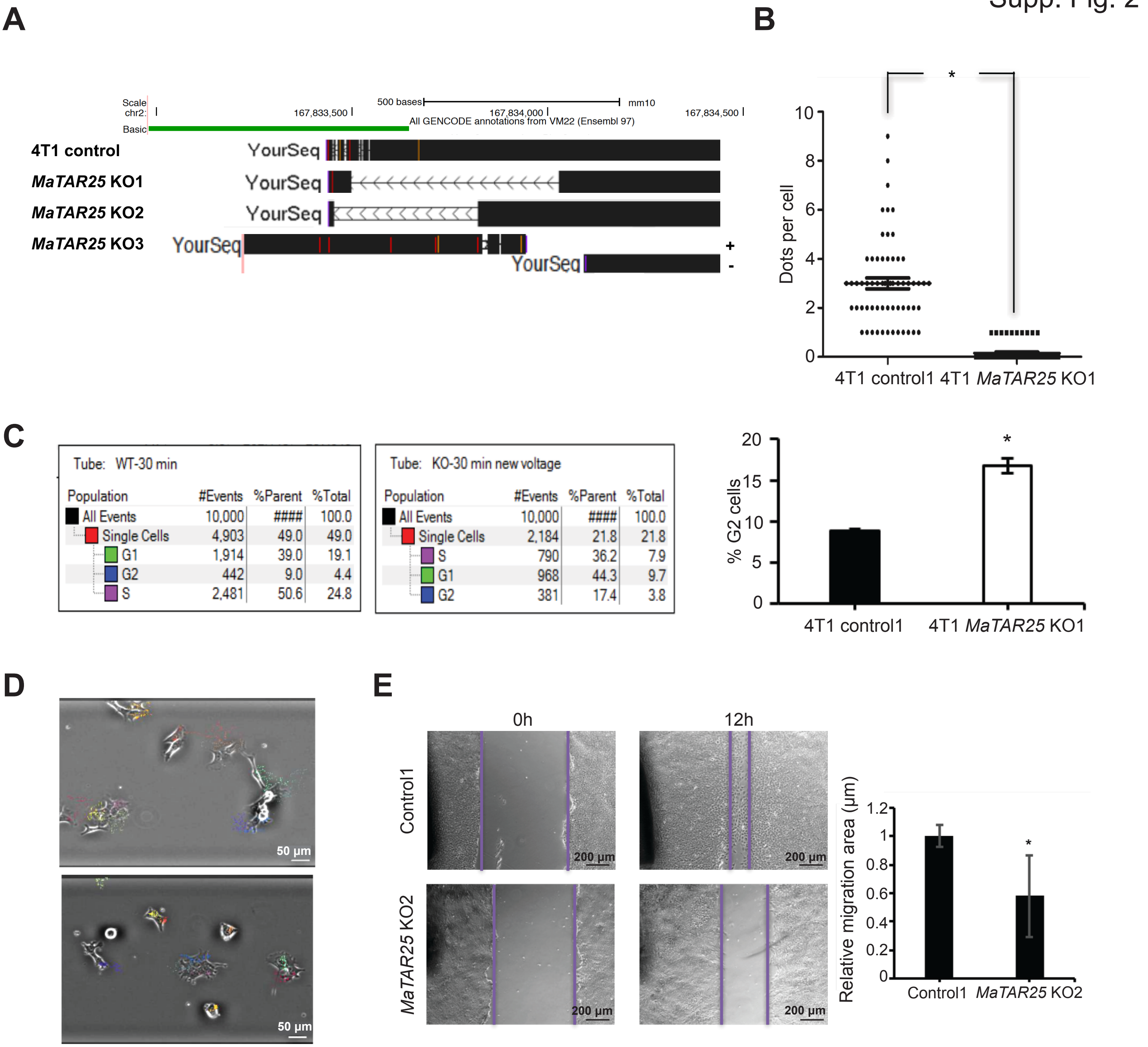


F

| | |
|---|---|
| 1 | DNA replication |
| 2 | Oxidative phosphorylation |
| 3 | Fanconi anemia pathway |
| 4 | Mismatch repair |
| 5 | Spliceosome |
| 6 | Pyrimidine metabolism |
| 7 | Glutathione metabolism |
| 8 | Steroid hormone biosynthesis |
| 9 | Proteasome |
| 10 | Homologous recombination |
| 11 | RNA transport |
| 12 | Cell cycle |
| 13 | Metabolism of xenobiotics by cytochrome P450 |
| 14 | Drug metabolism - cytochrome P450 |
| 15 | Fatty acid metabolism |
| 16 | Base excision repair |
| 17 | Ascorbate and aldarate metabolism |

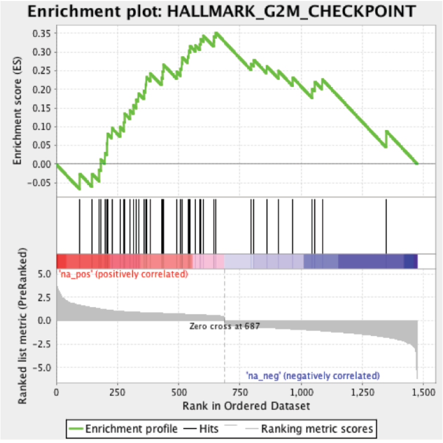


| | |
|---|---|
| Normalized Enrichment Score (NES) | 1.8573614 |
| Nominal p-value | 0.002631579 |
| FDR q-value | 0.022816334 |
| FWER p-Value | 0.093 |

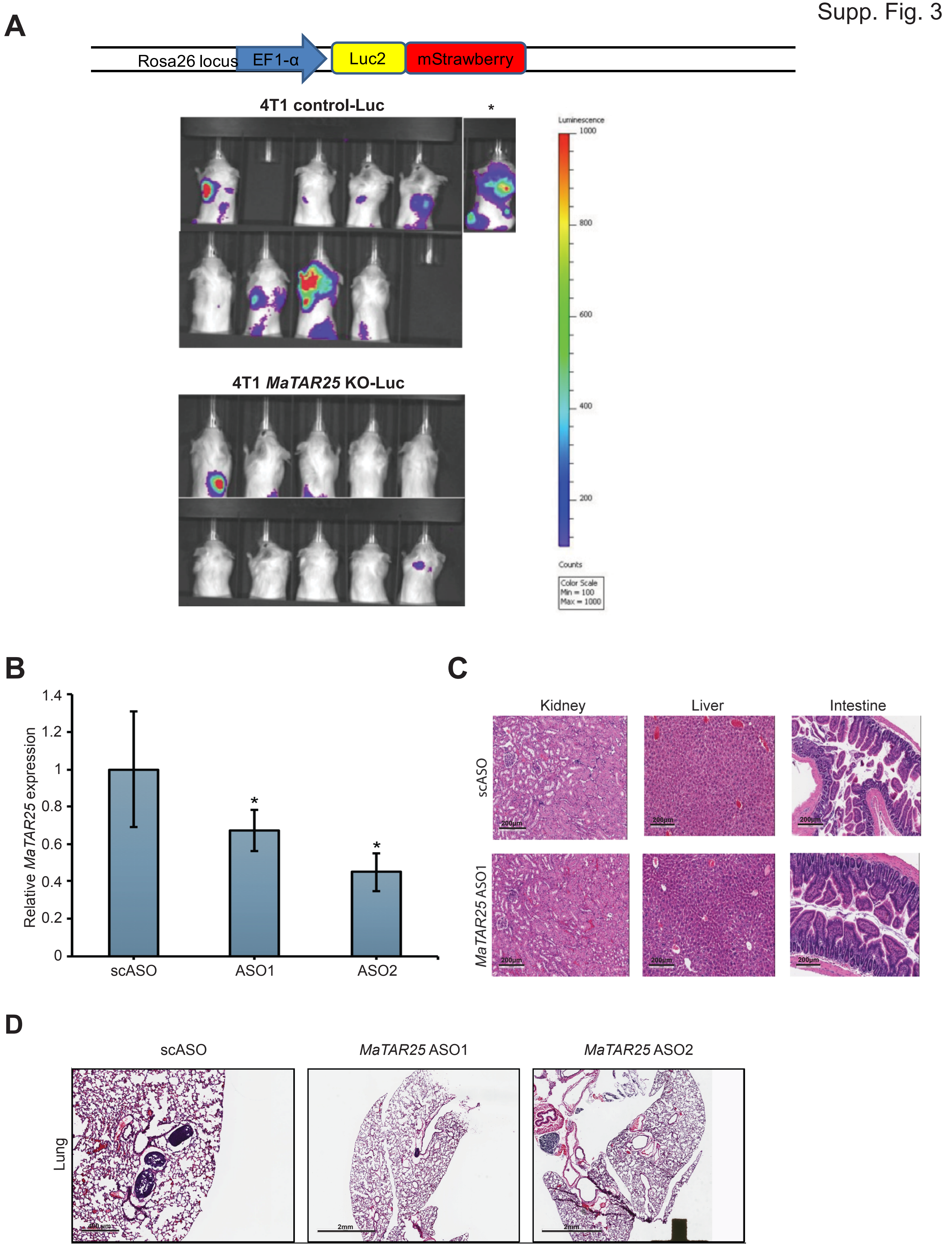
A
Rosa26 locus
EF1-α
Luc2
mStrawberry
4T1 control-Luc
*
4T1 MaTAR25 KO-Luc
Luminescence
1000
800
600
400
200
Counts
Color Scale
Min = 100
Max = 1000
B
Relative MaTAR25 expression
1.4
1.2
1
0.8
0.6
0.4
0.2
0
scASO
ASO1
ASO2
*
*
C
Kidney
Liver
Intestine
scASO
MaTAR25 ASO1
200μm
D
scASO
MaTAR25 ASO1
MaTAR25 ASO2
Lung
400μm
2mm
2mm

Supp. Fig. 4

A

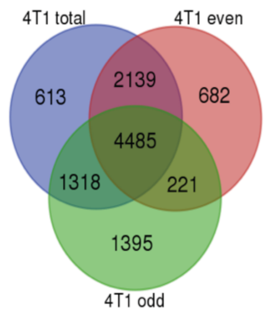


| Peak score | Detail annotation |
| --- | --- |
| 2778.6 | (TGG)n\|Simple_repeat\|Simple_repeat |
| 1771.9 | (CCA)n\|Simple_repeat\|Simple_repeat |
| 1164.2 | RLTR10C\|LTR\|ERVK |
| 1135.3 | CT-rich\|Low_complexity\|Low_complexity |
| 1091.7 | (TGG)n\|Simple_repeat\|Simple_repeat |
| 971.1 | (TGG)n\|Simple_repeat\|Simple_repeat |
| 963.2 | (TCCA)n\|Simple_repeat\|Simple_repeat |
| 872.4 | promoter-TSS (NM_023858) |
| 855.4 | ORR1B2\|LTR\|ERVL-MaLR |

B

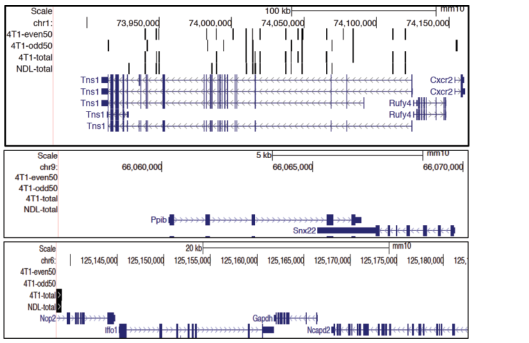

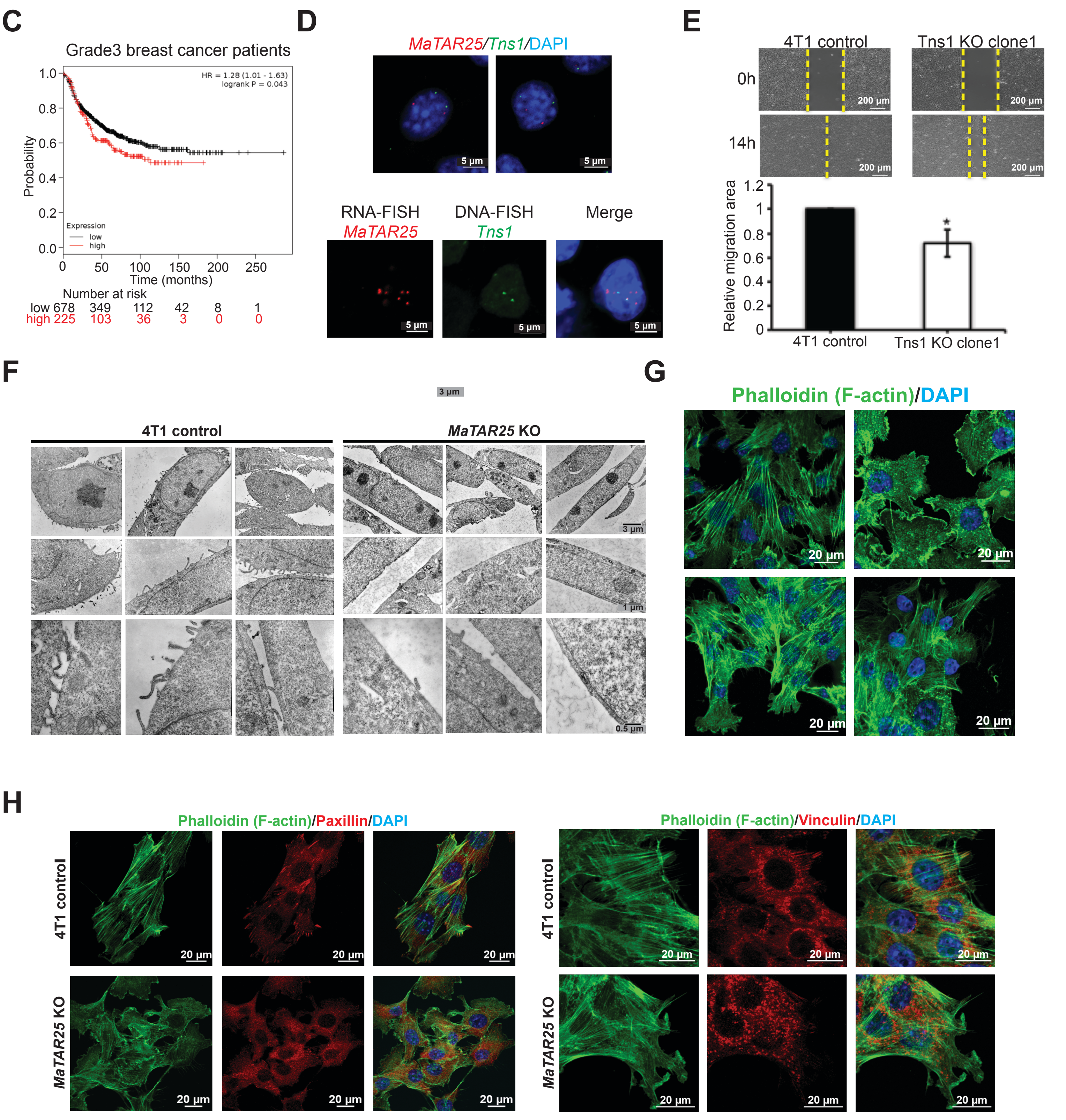

Supp. Fig. 5

**A**

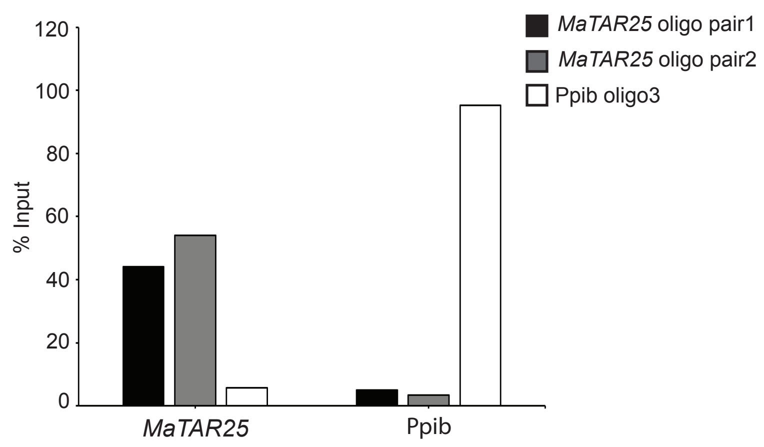


**B**

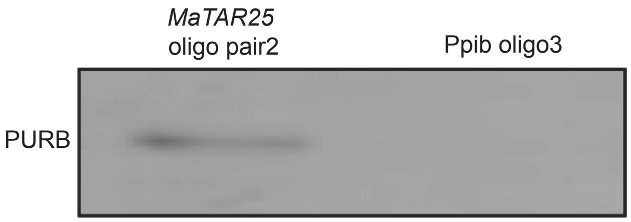


**C**

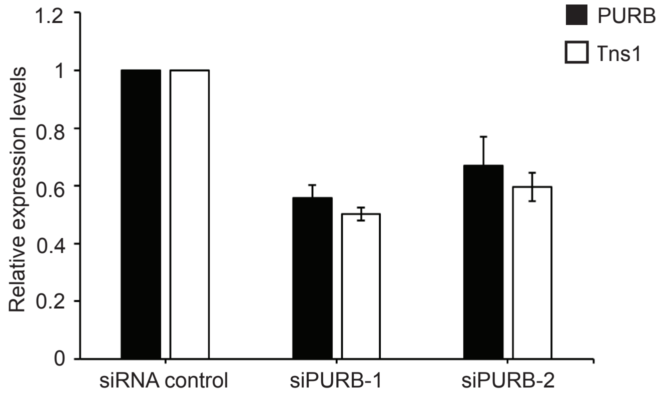


**D**

GAGTGATGAACATGATGGTGGAGATGATGGCG
ATGTAGGTGGTGGTGGTGGCGATGGAGGAGA
TATAGGTTATAGTGGTGATGACTGAGATGATGG
AGATGGGAATGGAGGTGGTGATGGAGATGAT
GCTGATGGAGGTGGTGGAGATGGGAATGGTA
GAGTTGGAGATGGTGGAGATGGAGGTGGTGA
AGATGGAGGTGGAGATGGTAGTGATGAAGAT
GGTAGTCACAGATAGTGGTGGCGGTGGTAATG

**E**

### Information for 1-GGTGGTGGAGAT (Motif 1)

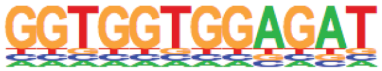


Reverse Opposite:

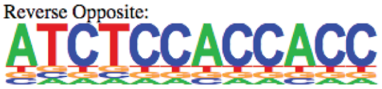


| | |
|---|---|
| p-value: | 1e-3288 |
| log p-value: | -7.571e+03 |
| Information Content per bp: | 1.530 |
| Number of Target Sequences with motif | 3991.0 |
| Percentage of Target Sequences with motif | 6.81% |
| Number of Background Sequences with motif | 243.0 |
| Percentage of Background Sequences with motif | 0.41% |
| Average Position of motif in Targets | 290.4 +/- 174.5bp |
| Average Position of motif in Background | 310.8 +/- 176.2bp |
| Strand Bias (log2 ratio + to - strand density) | -0.0 |
| Multiplicity (# of sites on avg that occur together) | 1.01 |
| Motif File: | file (matrix) reverse opposite |
| SVG Files for Logos: | forward logo reverse opposite |

Supp. Fig. 6

A

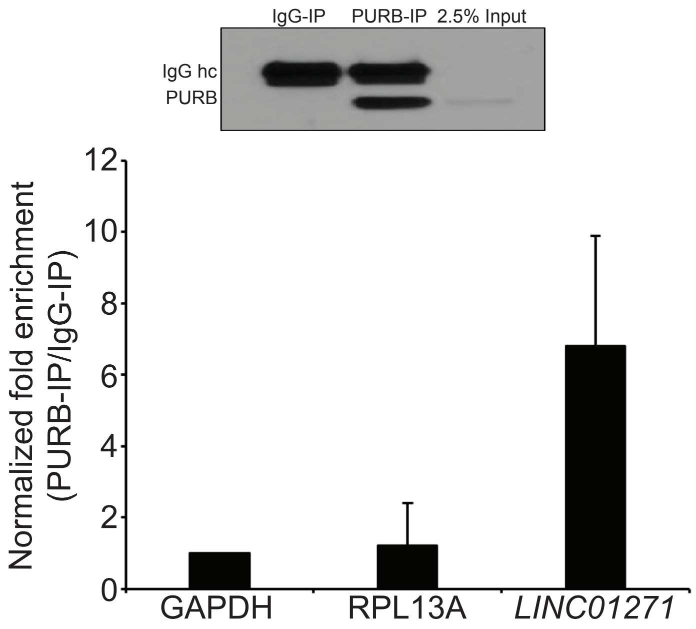


B

| Positive correclation |
|---|
| DNA REPLICATION |
| MITOTIC M M G1 PHASES |
| CELL CYCLE MITOTIC |
| M G1 TRANSITION |
| SYNTHESIS OF DNA |
| CELL CYCLE |
| S PHASE |

C

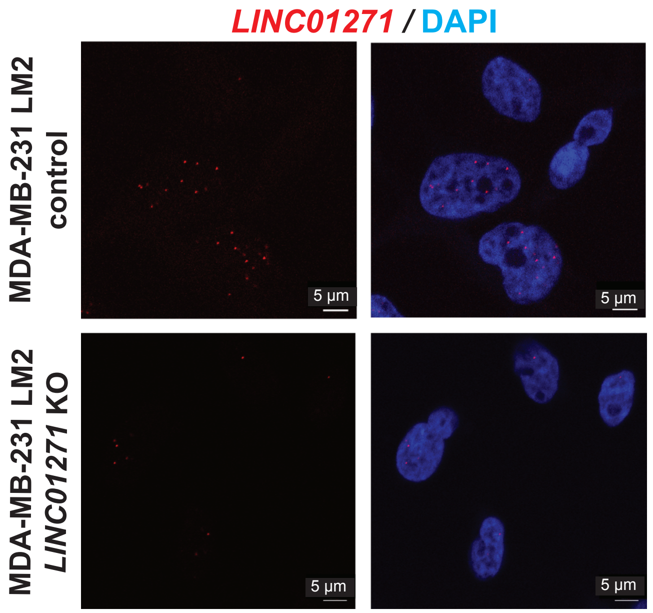


D

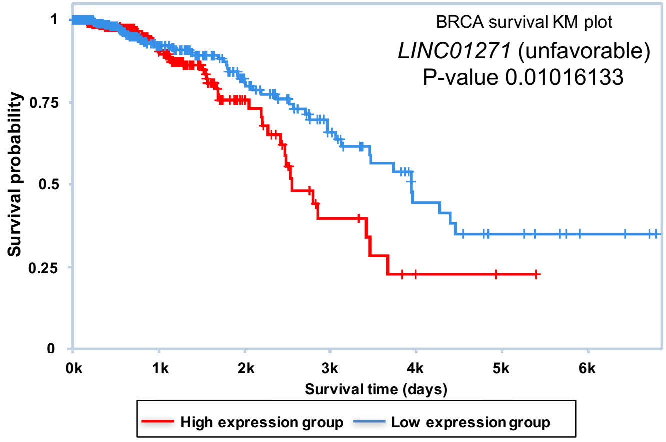


E

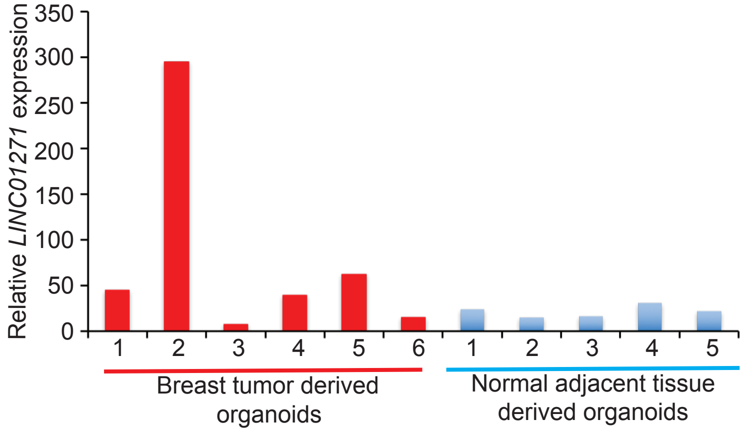

Supp. Fig. 7

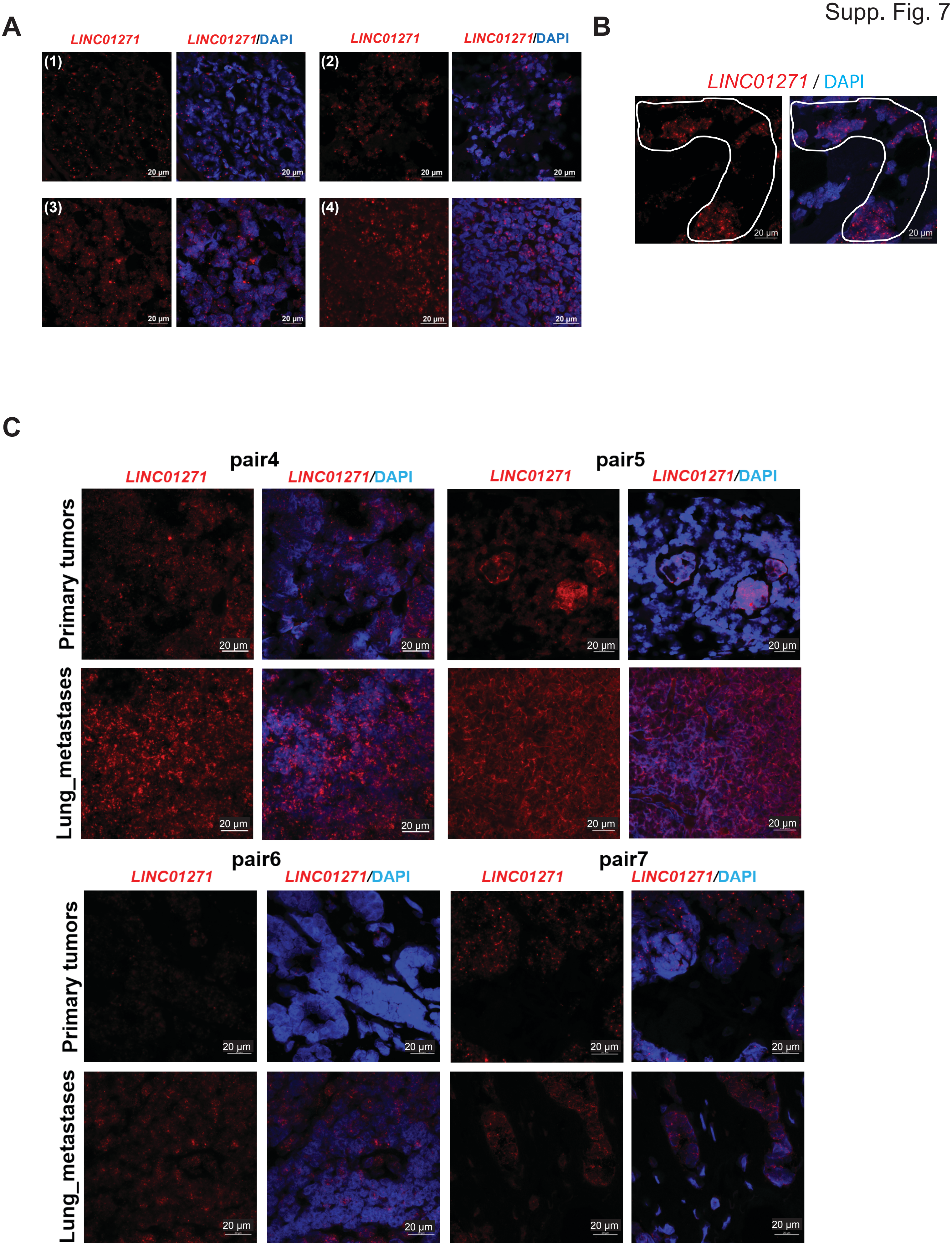
A
LINC01271
LINC01271/DAPI
(1)
(2)
(3)
(4)
20 μm
B
LINC01271 / DAPI
C
pair4
pair5
pair6
pair7
Primary tumors
Lung_metastases
