## Supplementary Table S3 for "*MaTAR25* LncRNA Regulates the *Tensin1* Gene to Impact Breast Cancer Progression"

| | Genome PCR for *MaTAR25* KO clone selection |
|---|---|
| Forward (F) | CCTGCCTCACACCACTTCTT |
| Reverse (R) | AGTTCACCTGTTGAGCAGAGC |

| | ChIP-PCR | Primer sequence |
|---|---|---|
| Murine | Tns1-outside of ChIRP identified region primer1-F | TGCTTCAGGTAGGGAGCTGT |
| Murine | Tns1-outside of ChIRP identified region primer1-R | CTATGCTGGCCTTGGCTTAG |
| Murine | Tns1-outside of ChIRP identified region primer2-F | CCATGTGGGTGATAGGAACC |
| Murine | Tns1-outside of ChIRP identified region primer2-R | CCTGGGTATCATCCCTAGCA |
| Murine | Tns1-inside of ChIRP identified region primer1-F | GTGGAGATGATGGCGATGT |
| Murine | Tns1-inside of ChIRP identified region primer1-R | CGCCACCACTATCTGTGACT |
| Murine | Tns1-inside of ChIRP identified region primer2-F | GGTGATGGAGATGATGCTGA |
| Murine | Tns1-inside of ChIRP identified region primer2-R | ATTACCACCGCCACCACTAT |
| Murine | Gapdh-promoter region-F | GAATGCCTTTTCTCCCTTCC |
| Murine | Gapdh-promoter region-R | GAGCCAGGGACTCTCCTTTT |

| | RT-PCR | Primer sequence |
|---|---|---|
| Murine | *MaTAR25* -primer1-F | TGATTTCCAACCTGGTCCCG |
| Murine | *MaTAR25* -primer1-R | ACTTAAGGTGGCCATGGCTC |
| Murine | *MaTAR25* -primer2-F | CCCTGCCTGGTTTCACTATG |
| Murine | *MaTAR25* -primer2-R | AGGCACAGGCTTTGGAAATA |
| Murine | *MaTAR25* -primer3-F | TGGGAAAAGGACAGACAACA |
| Murine | *MaTAR25* -primer3-R | CAGCCTCTGTCCCTGTCCT |
| Murine | *MaTAR25* -TSS-F | CCTGTGGAGCTTGGACAGTA |
| Murine | *MaTAR25* -TSS-R | GGGGGCTCTGGGTCTTTC |
| Murine | PPIB-F | GACAGACAGCCGGGACAAGC |
| Murine | PPIB-R | GGGGATTGACAGGACCCACA |
| Murine | PPIB-TSS-F | GAGCGCAATATGAAGGTGCT |
| Murine | PPIB-TSS-R | TGACTGTGACTTTAGGTCCCTTC |
| Murine | Tns1-primer1-F | AATGGAAAAGCTCCCTCTCC |
| Murine | Tns1-primer1-F | ATTTATGCCCTGCAACAAGG |

| | | |
|---|---|---|
| Murine | Tns1-primer2-F | TATGTCACCGAACGCATCAT |
| Murine | Tns1-primer2-R | AGGTGTCCATGGCCTTACAG |
| Murine | Tns1-primer3-F | ACTCCACCTAGCCCTGGTTT |
| Murine | Tns1-primer3-R | GGTTCCAGGAGTGGTAGCTG |
| Murine | PURB-F | ACAAGTACGGGGTGTTCCTG |
| Murine | PURB-R | TGTCGTCCTGGATCTCTTT |
| Murine | Ptpn1-F | TATTCTCACCCAGGGCCCTT |
| Murine | Ptpn1-R | TCTTCTTGCTGTGGCCAATA |
| Murine | Cebpb-F | ACTTCAGCCCCTACCTGGAG |
| Murine | Cebpb-R | GTAGTCGTCGGCGAAGAGGT |
| Murine | Snail1-F | ACGTCCGCACCCACACTGGT |
| Murine | Snail1-R | CTTCACATCCGAGTGGGTTT |
| Murine | Gapdh-F | GGTGGTGAAGCAGGCATCTG |
| Murine | Gapdh-R | CGGCATCGAAGGTGGAAGAG |
| Murine | Malat1-F | ACCAGTTTCCCCAGCTTTT |
| Murine | Malat1-R | CTACATTCCCACCCAGCACT |

| | | |
|---|---|---|
| Human | GAPDH-F | CGCTCTCTGCTCCTCCTGTT |
| Human | GAPDH-R | CCATGGTGTCTGAGCGAGT |
| Human | RPL13A-F | CCTGGAGGAAGAGGAAAGAGA |
| Human | RPL13A-R | TTGAGGACCTCTGTGTATTTGTCAA |
| Human | *LINC01271* -primer1-F | CAAGGGACAGAAAGCCAGTC |
| Human | *LINC01271* -primer1-R | TGACAGGAAGCAGCACAGAG |
| Human | *LINC01271* -primer2-F | GAATGGCCGAACTCACTTTC |
| Human | *LINC01271* -primer2-R | TAAGAGGTGGTTGGGTCCTG |
| Human | *LINC01270.1* -primer-F | CCAACGGTGTCCTTGTGTCT |
| Human | *LINC01270.1* -primer-R | CATCTGGGCTAACTGGTGCT |
| Human | *LINC01270.2* -primer-F | ACAGGAAGCACCAGTTAGCCCA |
| Human | *LINC01270.2* -primer-R | GGAAACTGAAGCCCAGAGAG |
| Human | TNS1-F | CTTTGCAAGGGGAAATCAAG |
| Human | TNS1-R | ACACACACCCACACGACAAT |

**CRISPR guide RNA sequences**

| | | gRNA sequence |
|---|---|---|
| Murine | *MaTAR25* TSS upstream gRNA1 | GTGGCCCCAGGGTGATGACT |
| Murine | *MaTAR25* TSS upstream gRNA2 | AACTTCCCGGCCTTATCGTC |
| Murine | *MaTAR25* TSS upstream gRNA3 | GCCTGACCTAGTTTCTCACC |
| Murine | *MaTAR25* TSS downstream gRNA | AAGTCAGACTCTTGGCGGGC |

| | | |
|---|---|---|
| Murine | *Tns1* gRNA1 | CGAGCGCCTCGCAGCACACC |
| Murine | *Tns1* gRNA2 | CTCACAGCGCTCGGGCCGCG |
| Murine | *Tns1* gRNA3 | GTGCGGCGCCCGACCCGAGG |
| Murine | *Tns1* gRNA4 | TGGTGGGGTTGACGGCCCGC |
| Murine | *Tns1* gRNA5 | ACGGCCCGCCGGCCAAAACC |

| | |
|---|---|
| Renilla luciferase control gRNA | GGTATAATACACCGCGCTAC |

**cEt ASO sequences**

| | | ASO sequence |
|---|---|---|
| Murine | *MaTAR25* ASO1 | GCATAGGAACTCAGGG |
| Murine | *MaTAR25* ASO2 | TTACATTCAGCTTTAG |
| Murine | *MaTAR25* ASO3 | TTTTTATTGGTATCAC |
| Human | *LINC01271* ASO1 | ATCTTTAGCAGAGGGA |
| Human | *LINC01271* ASO2 | GTCGGGTGGTCAAGGT |
| Human | *LINC01271* ASO3 | CCTACTGAAGAACCCC |

| | | |
|---|---|---|
| | scambled ASO (scASO) | CGCCGATAAGGTACAC |

**RACE Primer sequences**

| | | RACE specific primer sequence |
|---|---|---|
| *MaTAR25* | 5' RACE outer | CTACAGTCTGCCAGAACACTGC |
| *MaTAR25* | 5' RACE inner | TTCTGCTCGTGCTAGCCAAGT |
| *MaTAR25* | 3' RACE outer | GCCAAGACACAGAGAACACG |
| *MaTAR25* | 3' RACE inner | GTACATACCCAGTGCCCTGA |

**Biotin probe sequences**

| | 3' Biotin-labelling oligonecleutide probe sequence |
|---|---|
| *MaTAR25*- oligo1 | GGATCAATACTGTCCAAGCT |
| *MaTAR25*- oligo2 | AAGAGTCTGACTTCTTGGCA |
| *MaTAR25*- oligo3 | TCCTGGAGATGCATTTTTTC |
| *MaTAR25*- oligo4 | CAGGTGGTGGAGATGGAATG |
| *MaTAR25*- oligo5 | GAAAGGAAACACCCAGGTCA |
| *MaTAR25*- oligo6 | AGTCACAGAGAAGGACGGCG |
| *MaTAR25*- oligo7 | TTGGGAGTAACCTGGGGAAG |
| *MaTAR25*- oligo8 | CTTGCTCCAGAACGACAGCG |
| *MaTAR25*- oligo9 | CAGTCACACACTTGACACAT |
| *MaTAR25*- oligo10 | ATGATACCTCTCATCTGGAG |
| *MaTAR25*- oligo11 | AGAAGGAATGTCACCTGTCA |
| *MaTAR25*- oligo12 | AATAGGGAGAAGGGAGGCTC |
| *MaTAR25*- oligo13 | CTCAATTCACCCATGGATTA |
| *MaTAR25*- oligo14 | GGCTTTGGAAATACATTGGG |

| | |
|---|---|
| Ppib-oligo1 | GACCGGAAAAGGGTGTGGAC |
| Ppib-oligo2 | CGGGCAGCAAAAGGAAGACG |
| Ppib-oligo3 | CCTACAGATTCATCTCCAAT |

| | |
|---|---|
| Ppib-oligo4 | GAAGTTCTCATCTGGGAAGC |
| Ppib-oligo5 | GTTATGAAGAACTGTGAGCC |
| Ppib-oligo6 | CCGGAGTCGACAATGATGAC |
| Ppib-oligo7 | ACACAGACAGCTGCTTAGAG |
| Ppib-oligo8 | AAATCAGGCCTGTGGAATGT |
| ***RNA Antisense Pulldown and Mass Spectrometry** | |
| oligo pair1<br>oligo pair2 | *MaTAR25* -oligo7 and oligo8<br>*MaTAR25* -oligo6 and oligo10 |

| | **Primer sequence** |
|---|---|
| MMTV-Neu-NDL genotyping-F | TTCCGGAACCCACATCAGGCC |
| MMTV-Neu-NDL genotyping-R | GTTTCCTGCAGCAGCCTACGC |
