## Supplementary Figure Legends for "*MaTAR25* LncRNA Regulates the *Tensin1* Gene to Impact Breast Cancer Progression"

**Figure S1. Characterization of *Mammary Tumor Associated RNA 25* (*MaTAR25*).**

**Related to Figure 1.**

(A) Expression profile of *MaTAR25* in various mouse tissue types from ENCODE data sets compared to RNA-Seq results of mammary tumor from MMTV-Neu-NDL mice (Diermeier et al., 2016), and from FANTOM5 database (Petryszak et al., 2016).
(B) Screen shot of *MaTAR25* genomic locus from UCSC genome browser showing PhyloCSFraw scores of individual codons in each of three frames (+ and – strands; horizontal line is 0).
(C) Protein coding potential analysis of *MaTAR25* RNA transcript using Coding-Potential Assessment Tool (CPAT).
(D) Assessing the protein coding potential of *MaTAR25* using Coding Potential Calculator (CPC), and the predicted open reading frame (ORF) and putative peptide.
(E) qRT-PCR showing the knockdown efficiency of 2 independent ASOs targeting *MaTAR25* in 4T1 cells after 48 hours and 72 hours of incubation compared to mock and scASO treated control cells. The result is shown ± SD (n=3).
(F) 4T1 cells were seeded at the same cell density (5x10^4^/well) in 12-well tissue culture plates at day 0 while ASOs were added into the culture medium and cell counting was performed at different time points to measure cell numbers. The mean cell numbers of three independent replicates of 4T1 mock treated control cells, 4T1 cells treated with scrambled ASO, 4T1 cells treated with 2 different *MaTAR25* ASOs is shown ± SD (n=3) **p* < 0.05 (student’s t-test).

**Figure S2. *MaTAR25* knockout affects 4T1 cell cycle progression, and migration as well as related regulatory pathways *in vitro.* Related to Figure 2.**

(A) A snapshot of BLAT results showing the CRISPR-Cas9 mediated editing events of 3 different 4T1 *MaTAR25* knockout (KO) clones over the promoter region of the *MaTAR25* gene locus by aligning Sanger sequencing results on the UCSC genome browser.
(B) Quantitation of the number of *MaTAR25* smRNA-FISH foci in individual 4T1 control and *MaTAR25* KO cells. Over 100 cells were counted in each group and the result is shown ± SD **p* < 0.05 (student’s t-test).
(C) BrdU incorporation cell cycle analysis using flow cytometry to compare the percentage of cells in each cell cycle stage between 4T1 control and *MaTAR25* KO cells. The percentage of cells in G2 phase in 4T1 control vs *MaTAR25* KO cells ± SD (n=2) **p* < 0.05 (student’s t-test).
(D) Representative live cell images from tracking 4T1 control cells and *MaTAR25* KO cells over time.
(E) Scratch wound healing assay showing the difference in migration ability between 4T1 control vs *MaTAR25* KO cells. The wound line was created by a microtip, then cells were rinsed with PBS twice before adding new medium and incubated for 12 hours. The migration areas in each group were measured by ImageJ and the result is shown ± SD (n=2) **p* < 0.05 (student’s t-test).
(F) Pathway analysis showing affected pathways in *MaTAR25* KO cells when comparing differentially expressed genes identified by RNA-Seq analysis from *MaTAR25* KO cells compared to 4T1 control cells.

**Figure S3. *MaTAR25* knockout impairs lung metastasis *in vivo.* Related to Figure 3.**

(A) Schematic representation of the plasmid construct targeted into the Rosa26 locus for positive clone selection and *in vivo* bioluminescence imaging of luciferase labeled 4T1 control and *MaTAR25* KO cells. The *in vivo* bioluminescence images were acquired at day 21 from female BALB/c mice tail vein injected with 4T1 control or *MaTAR25* KO cells.
(B) qRT-PCR analysis showing the *in vivo* knockdown efficiency of 2 independent ASOs targeting *MaTAR25* in tumors collected from MMTV-Neu-NDL mice compared to tumors from scASO injected mice. The result is shown ± SD (n=7) **p* < 0.05 (student’s t-test).
(C) Representative hematoxylin and eosin (H&E) stained tissue images (kidney, liver and intestine) showing no histological phenotypes between scASO treated and *MaTAR25* ASO treated groups. Normal tissues are not impacted by *MaTAR25* knockdown.
(D) Representative hematoxylin and eosin (H&E) stained lung images showing reduced micro-metastatic nodules in lung tissue from mice treated with each of two *MaTAR25* ASOs vs scASO control.

**Figure S4. *MaTAR25* binds to the *Tns1* gene body, and *Tns1* is the key downstream target of *MaTAR25* for regulating tumor cell progression and migration*.* Related to Figure 4.**

(A) Venn diagram showing the overlap of genomic target sites from ChIRP-Seq data using *MaTAR25* total oligo pool (14 oligos), *MaTAR25* odd oligo pool (7 oligos), or *MaTAR25* even oligo pool (7 oligos) for RNA pull-down in 4T1 cells. The number of target sites is indicated by the overlap between the circles.

Top genome ontology enrichment regions from analyzing the sequence of overlapped target sites are listed in the right table.

(B) UCSC genome browser screen shot showing peaks of *MaTAR25* ChIRP-Seq data over *Tns1* and two non-target genes (*Ppib, Gapdh*) using *MaTAR25* odd oligo pool (7 oligos), *MaTAR25* even oligo pool (7 oligos), or *MaTAR25* total oligo pool (14 oligos) for RNA pull-down in 4T1 cells, and *MaTAR25* total oligo pool (14 oligos) for RNA pull-down in MMTV-NDL cells.
(C) Kaplan Meier-plotter analysis by KM plotter (Gyorffy et al., 2010) showing the correlation of *Tns1* expression levels (lower and upper quartiles were computed and indicated as low expression and high expression groups) with survival of patients with Grade 3 breast tumors.
(D) Representative images of double DNA-FISH detecting *MaTAR25* and *Tns1* gene loci within 4T1 cells (upper). Representative images of smRNA-FISH of *MaTAR25* RNA transcripts and DNA-FISH detecting *TNS1* gene loci within the same cell to assess the potential interaction between *MaTAR25* RNA transcripts and *Tns1* gene loci in 4T1 cells (lower).
(E) Scratch wound healing assay showing the difference in migration ability between 4T1 control and *Tns1* KO cells. The migration areas in each group were measured by ImageJ and the result is shown ± SD (n=2) **p* < 0.05 (student’s t-test).
(F) Representative transmission electron microscopy (TEM) images showing the significant reduction in microvilli upon MaTAR25 KO. Shared scale bars are indicated in the right panels.
(G) Representative images showing the changes of F-actin bundles in (1) 4T1 control cells (2) 4T1 *MaTAR25* KO cells (3) 4T1 *MaTAR25* KO cells with ectopic expression of *MaTAR25* (4) 4T1 *MaTAR25* KO cells with ectopic expression of Tns1. Cells were stained with Alexa Fluor 488 phalloidin conjugate, and images were acquired using a Zeiss 710 confocal microscope. Scale bars are 20 μm.
(H) Representative images showing the changes in the localization of focal adhesion complex proteins in 4T1 control and 4T1 *MaTAR25* KO cells by immunofluorescence (IF) labeling of paxillin (upper) and vinculin (lower). Cells were co-stained with Alexa Fluor 488 phalloidin conjugate. Scale bars are 20 μm.

**Figure S5. *MaTAR25* interacts with PURB to carry out its function. Related to Figure 5.**

(A) *MaTAR25* or PPIB (control) RNA transcripts were assessed by qRT-PCR following the pull-down of *MaTAR2*5 or *PPIB* in 4T1 cells.
(B) Immunoblot analysis of PURB following the pull-down of *MaTAR25* or PPIB in MMTV-Neu-NDL primary cells.
(C) qRT-PCR analysis of *Tns1* expression in 4T1 cells following treatment for 48 hours with three independent siRNAs to knockdown PURB, compared to negative control cells.
(D) The DNA targeting sequence of *Tns1* DNA recognized by *MaTAR25* by ChIRP-Seq analysis. The PURB binding sequence motif (GGTGG) is highlighted.
(E) An enriched transcription factor motif sequence was identified from *MaTAR25* ChIRP-seq using HOMER (Heinz et al., 2010 ).

**Figure S6. *LINC01271* is the human ortholog of *MaTAR25*. Related to Figure 6.**

(A) RNA immunoprecipitation (RIP) and qRT-PCR using PURB antibody to identify the specific interaction between PURB and *LINC01271* RNA transcripts in MDA-MB-231 LM2 cells. Rabbit IgG antibody was used as a negative control. qRT-PCR detecting *Gapdh* and *Rpl13a* RNA transcripts were used as non-specific RNA transcript binding for normalization.
(B) Identification of the positive correlation pathways with *LINC01271* expression by reactome analysis based on the Breast Cancer TCGA data.
(C) Representative images of smRNA-FISH detecting *LINC01271* RNA transcripts in MDA-MB-231 LM2 cells to assess the localization of *LINC01271* in control and *LINC01271* KO cells. Scale bars are 5 μm.
(D) Kaplan Meier-plotter analysis by TANRIC (Li et al., 2015) showing the correlation of *LINC01271* expression level (lower and upper quartiles were computed and indicated as low expression and high expression groups) on survival of breast cancer patients.
(E) Comparison of the expression level of *LINC01271* within tumor organoids derived from samples of luminal subtype breast cancer patient vs normal breast organoids derived from adjacent breast tissues detected by qRT-PCR.

**Figure S7. smRNA-FISH examining *LINC01271* expression in patient breast tumor samples. Related to Figure 7.**

(A) Representative smRNA-FISH images showing the expression of *LINC01271* in patient breast tumor sections from early stage (1) (2) and late stage (3) (4) breast tumor sections. Scale bars are 20 μm.
(B) Representative smRNA-FISH images showing the clonal/regional expression pattern of *LINC01271* RNA transcripts within breast tumor patient samples. Cells with higher expression level of *LINC01271* are highlighted. Scale bars are 20 μm.
(C) smRNA-FISH images showing the expression pattern of *LINC01271* within luminal subtype breast cancer primary tumors and lung metastases sections from the same patients. Scale bars are 20 μm.
